# Ecological genomics of preference for human hosts in the native range of the dengue mosquito

**DOI:** 10.64898/2026.02.11.705442

**Authors:** James E. Fifer, Massamba Sylla, Mathias Ouedraogo, Abdoul-Aziz M. Maiga, Aboubacar Sombié, Boubacar Coulibaly, Binta Djimde, Sanou Makan Konate, Karim Sawadogo, Sale Sidibe, Sakina Isadibir, Dvorah Nelson, Yunusa M. Garba, Athanase Badolo, Alpha Seydou Yaro, Noah H. Rose

## Abstract

The arbovirus transmitting mosquito *Aedes aegypti* has African origins where it maintains generalist and human specialist feeding forms, the latter of which has recently expanded their range worldwide. These two different forms appear to be directly linked to discrete genetic lineages, where the relative proportion of each ancestry dictates a mosquito’s standing on the human specialist-generalist behavior spectrum. Here we take a population genomics approach to develop a predictive model for describing the distribution of human specialist behavior across Africa. We show genome-wide ancestry is predictive of human specialist behavior, but a subset of the genome offers even greater predictability. We also show that an ecological model incorporating precipitation, temperature and human density explains the geographic patterning of human specialist behavior in Africa.

## Introduction

The mosquito *Aedes aegypti*, the primary vector of dengue, yellow fever, Zika and chikungunya, is one of the largest contributors to the emergence and spread of vector-borne disease (Powell et al., 2018). The outsized contribution of *Ae. aegypti* to the vector-borne disease burden is due to its recently evolved adaptations to seeking human hosts and living in human habitats, allowing for exploitation of anthropogenic environmental niches. In Africa, the native range of *Ae. aegypti*, this mosquito is found as two distinct forms: a generalist that lives in both forests and rural human settlements, *Ae. aegypti formosus (Aaf)*, and the human-specialist *Ae. aegypti aegypti (Aaa)*, the latter of which dominates the invasive range. Coinciding with a rising burden of *Aedes-transmitted* (Selhorst et al., 2020; Willcox et al., 2018) - outbreaks of dengue in 15 of the 47 African countries (World Health Organization, 2023) and potential endemicity in at least 34 African nations (Gainor et al., 2022)- behavioral variation across *Ae. aegypti’s* African range positions the continent as an ideal setting for ecological genomics work on host preference, the trait that underlies much of the species’ exceptional ability to serve as an effective vector in human epidemics (Caldwell et al., 2024).

Previous studies investigating the ecological and genomic drivers of contemporary human preferring populations in Africa revealed the presence of multiple genetic lineages of *Ae. aegypti* (Brown et al., 2011; Crawford et al., 2017; Gloria-Soria et al., 2016; Kotsakiozi et al., 2018; Moore et al., 2013; Mulwa et al., 2025; Rose et al., 2020; Tabachnick & Powell, 1978) and that relative proportions of respective ancestries dictate where a mosquito falls on the human-specialist to generalist host preference spectrum (Rose et al., 2020). A first attempt at building an ecological model indicated that human population density and a climate with high precipitation seasonality may play large roles in shaping how human host preference is partitioned across *Ae. aegypti’s* native range (Rose et al., 2020). The Sahel region of Western Africa, in particular, exhibits high precipitation seasonality as well as containing populations with a high proportion of human-specialist ancestry and high human preference. The long dry seasons produced by high precipitation seasonality likely increases the dependence of *Ae. aegypti* on human stored water to complete their life cycle, which in turn favors an ecotype better adapted to feeding on the most abundant nearby host, humans (Rose et al., 2023).

An ecological genomics approach to explore host preference variation in *Ae. aegypti* could provide useful tools and insights for managing arboviral disease. Working towards a genomic predictor of preference can help identify candidate causal genes for functional studies and predictive markers that reduce reliance on labor-intensive behavioral assays when estimating vectorial capacity. Ecological models predicting the distribution of human preferring mosquito populations can assist with vector control efforts, for example the preliminary ecological model from Rose et al (2020) showed the ability to predict ZIKV seroprevalence variation in Africa (Caldwell et al., 2024). However, despite the promise of this approach, there are still several knowledge gaps in this system that are limiting inferences. Initial work was based on a small number of populations and had limited sampling in the Sahel region, which showed the most sharply variable patterns of host preference. Here we address several unanswered questions from this system. We sampled heavily in the Sahel region to validate the relationship between host preference and genetic ancestry, find regions of the genome that are most important for predicting host preference, and then build ecological models for predicting contemporary distributions of human-preferring populations of *Ae. aegypti* in Africa.

## Methods

### Field collections

A first round of collections was carried out in 2017 and 2018, with the exception of Uganda collections from 2015, and was discussed in previously published work (Rose et al., 2020). A second round of collections were obtained in the 2020 rainy season. These collections consisted exclusively of larvae, in order to maximize geographic breadth. Samples were collected from tires, watering cans and plastic bottles. One larva per breeding site was used downstream. A third round of collections were obtained in the 2021 rainy season using the same methods as described by Rose et al 2020 for the 2017 and 2018 collections. To briefly summarize, mosquito eggs were collected by distributing 20-60 ovitraps across the landscape and retrieving them after three nights. Eggs were dried on seed papers for 24 h and brought back to the laboratory where eggs were hatched and reared to adults and then either used for behavior or frozen for DNA extractions. All sampling information is in Supplemental Table 1.

### Behavior

We tested the host odor preference 7- to 14-day-old females following the methods from Rose et al 2020. This involves placing mosquitoes in a two-port olfactometer where one host chamber contains an awake guinea pig (*Cavia porcellus*, pigmented breed) and the second chamber contains a section of the arm of a human volunteer. All human trials used a 31-year-old European-American male and two guinea pigs were alternated. Three colonies from Rose et al 2020 (ORL, NGO and U52) were included as controls to ensure 2021 and 2017/2018 datasets were comparable. Odor preference for each population was summarized following Rose et al (2020), briefly-a beta-binomial mixed generalized model, with a fixed effect of population and random effects of day and colony, was used via the R package glmmTMB to model the population’s probability of choosing a human versus animal host. We ran two colonies, one human specialist (NGO) and one generalist (U52), across both timepoints to control for possible batch effects (Supp. Fig. 16). The fitted probability (p) of choosing a human host was transformed into a preference index (2p-1) where an index of zero means mosquitoes from that population were equally likely to choose either host.

### Whole Genome Sequencing (WGS)

This study combined several different datasets (see Supp. Table 1 for further breakdown) of *Ae. aegypti*: (i) ∼10-15x coverage WGS dataset of adult female and male *Ae. aegypti* from the 1206 *Ae. aegypti* genomes project, we used all 390 individuals from the 32 African populations as well as 20 individuals from Bangkok, Thailand and 4 Ae. mascarensis mosquitoes for use as an outgroup (Crawford et al., 2025). (ii) 1-2x coverage WGS of adult female and male *Ae. aegypti* from the 2021 collections which included 109 individuals from 12 African Sahelian populations. (iii) 1-2x coverage WGS of larval *Ae. aegypti* from 2020 rainy season which included 55 individuals from 18 populations in Senegal. (iv) ∼10-15x coverage WGS dataset of 15 adult female and male *Ae. aegypti* from Cape Verde (Rose et al., 2022). For (ii) and (iii) we extracted DNA using the NucleoMag 96 well extraction and tagmentation-based whole genome libraries for low coverage sequencing adapting protocols from (Langdon et al., 2022). Libraries were sequenced using NovaSeq XPlus 150PE.

All sequences were mapped to a version of the L5 reference genome (Matthews et al., 2018) that was updated with WGS data from 100 unrelated African male *Ae. aegypti* (Rose et al., 2020). Reads were trimmed using TRIMMOMATIC [ILLUMINACLIP:NexteraPE-PE.fa:2:30:10:2:keepBothReads LEADING:3 TRAILING:3 MINLEN:36] mapped to the reference using bwa (Li & Durbin, 2009) converted to bams and filtered to only include reads with mapping quality >20 with samtools (Li et al., 2009), duplicates removed using picard (“Picard Toolkit,” 2019) and clipping overlaps with BamUtil’s clipOverlap (Jun et al., 2015).

ANGSD (Korneliussen et al., 2014) was used for all genotype calling. For likelihood-based analyses (FastNGSadmix, NGSadmix) we used filters -uniqueOnly 1 -remove_bads 1 -minMapQ 20 -minQ 25 -dosnpstat 1 -doHWE 1 -skipTriallelic 1 -setMaxDepthInd 15 -minMaf 0.01 here. We also created allele frequency files for several analyses [f4-ratio, LFMM, ngsRelate] using likelihood-based calling methods with filters -minMapQ 10 -minQ 20 using a previously identified set 14,045,728 high confidence SNP calls (Rose et al., 2020) for major and minor allele classifications. Related individuals were filtered out using ngsRelate (Korneliussen & Moltke, 2015), using the coefficient of kinship estimate to remove putative first cousins (theta = 0.0625).

### Population structure analyses to predict behavior

We used FastNGSadmix (Jørsboe et al., 2017) to estimate each population’s assignment to the three major genetic ancestry groups present in *Ae. aegypti*: *Ae. aegypti* formosus East (AafE), *Ae. aegypti* formosus West (*AafW*) and *Aaa* (*i*.*e*. human-specialist ancestry). To estimate the proportions of these three ancestries for each sample we created a reference panel from the 1206 *Ae. aegypti* genome project with unadmixed samples from each of the three major ancestry groups (21 individuals with unadmixed *AafE*, 47 individuals with unadmixed *AafW* and 455 unadmixed *Aaa*). The loci used was a set of previously bcftools-called biallelic SNPs (Rose et al., 2020; 2023) the existing filters were as follows: covered by at least 1 read in 90% of individuals and then called for any individual with sample depth > 8 reads and genotype quality score > 30. After individual genotype calling sites were further filtered for the fraction of individuals genotyped (> 75% at any given SNP), minor allele frequency (MAF > 1%), MAPQ cutoff of 10 and GQ >20 in putative single-copy regions and excluding centromeric and repeat regions and the sex locus (Matthews et al., 2018). The panel consisted of allele frequencies for each of these three refs at 18,434,131 loci. While the dominance of these three ancestry groupings has been demonstrated in several African range population genomic studies of *Ae. aegypti* (Crawford et al., 2017; Kotsakiozi et al., 2018; Rose et al., 2020) we also used NGSadmix (Skotte et al., 2013) to confirm that K of 3 was meaningful for our dataset. We ran NGSadmix for Ks 1-8 with four replicates per each K and used the Evanno method (Evanno et al., 2005) for selecting K (Supp. Fig. 2). Also to ensure that differences in coverage were not biasing estimates from FastNGSadmix we downsampled 60 of the ∼10-15x samples to 2x, 1x, 0.1x, 0.001x and 0.0001x randomly selecting loci with 5 reps at each depth and then re-estimated population assignment with FastNGSadmix (Supp. Fig. 1). Genome-wide human-specialist ancestry proportions were compared to host preference data using a generalized additive model (GAM) via the R (R Core Team, 2024) package mgcv (Wood, 2017) with a beta regression family and with the shrinkage smoother “ts” to allow for non-linear associations between ancestry and our behavioral data. We rescaled preference to be between 0 and 1 ((Preference + 1) / 2) to deal with data frequently approaching -1 and accommodate beta regression assumptions.

We used windowed f4-ratio tests (10mb window, 10kb step) to identify regions of the genome where human-specialist ancestry was particularly predictive of preferring human hosts. We randomly split samples from Bangkok, a population that represents one of the most derived lineages of *Ae. aegypti* (Crawford et al., 2025), to create pseudo independent lineages (hereafter BKKa and BKKb) following (Malinsky et al., 2021). To avoid violating topological assumptions of f4-ratio while managing admixture from different Ae. ae. formosus lineages (i.e. *AafE* or *AafW*), we use two different trees depending on the presence of *AafE*. For populations with no AafE, we used a tree with OHI (unadmixed *AafW*), BKKa, query population, BKKb with SHM (unadmixed AafE) as the outgroup (Fig. 2B). For populations with admixture from *AafE* we instead use a tree with SHM (unadmixed *AafE*), BKKa, query population, BKKb with *Ae. masc* as an outgroup (Fig. 2C). We use a two-tree approach instead of a single-tree because while it is possible to also use the latter tree to estimate f4-ratio in populations with *AafW*, this would run the risk of near *Aaa AafW* being conflated with *Aaa*.

To find the most predictive windows we ran separate GAMs for each 10mb window and iterated across R^2^ percentile thresholds (0.60-0.999, increasing by 0.001). At each threshold the predictive power of all windows passing that threshold was assessed with a second GAM. We repeated this analysis using a LOOCV cross validation and also adding genome-wide *Aaa* ancestry as a covariate with the goal of seeing whether any windows had an outsized impact when controlling for the genome-wide signal. The threshold that maximally predicted preference (per R^2^) was selected for downstream host preference predictions. We also used GAM model selection to investigate whether a particular region (defined as consecutive 10mb windows that passed our best threshold) was more predictive than others and if different regions have discrete, non-linear effects. For the predictive window analysis, we used GAMs with logit transformed data and a Gaussian family instead of a beta regression family. We switched to this method because we wanted to use these predictive windows to predict preference for populations without behavior data downstream, and the no behavior populations included a subset with ancestry outside the range of ancestry inputted in our models. GAMs with the beta regression family can cause issues in this specific circumstance when best fitting edfs are > 1, leading to predictions outside of the possible parameter space (*i*.*e*. out the preference index of -1 to 1), a logit transformation with a Gaussian family GAM circumvents this issue.

To assess whether single loci were predictive of preference we leveraged latent factor mixed models (LFMMs) which allow us to control for the genome-wide signal of ancestry-preference associations that is present in our data. We ran LFMMs with K of 3 and genomic.control=TRUE using the R package LEA (Gain & François, 2021). We used an FDR correction to identify outliers.

### Ecological models

We used GAMs to find the environmental variables that best predict contemporary human host preferring distributions in two ways; using either real host preference or predicted host preference from the candidate regions identified with the f4-ratio analyses. We included all bioclim2 and CHELSA V2.1 variables (Karger et al., 2021) as predictors as well as contemporary (2015) farming practices (Dixon et al., 2020) and urban density. For CHELSA V2.1 data we excluded all “growing degree days” and “snow cover days” because they did not vary between our locations. For contemporary farming practices we tested the predictability of agropastoral farming practice dominance for host preference in several different ways, choosing the most informative method of summarizing the data with beta regressions. We coded agropastoral practice as a binary variable (i.e. either the area’s dominant practice was agropastoral farming or it was not) and also looked at the proportion of the dominated by agropastoral farming using 5 different buffer regions (5km, 10km, 20km, 50km, 100km). We rescaled preference to be between 0 and 1 ((Preference + 1) / 2) to deal with data frequently approaching -1 and accommodate beta regression assumptions. In a beta regression model that also takes regional variation into account (Preference = agropastoral_buffer + Latitude*Longitude), 100km explained the most variation. We also used a 20km buffer generated from a 2.5-min resolution population density raster from the 2015 United Nations World Population Prospects (United Nations, Department of Economic and Social Affairs, Population Division, 2015; Warszawski et al., 2017), for urban density. This buffer size was selected based on its predictability of host preference from Rose et al (2020).

GAM model selection was performed using the beta regression family with the shrinkage smoother “ts” as this smoother is well equipped for modelling spatial data (Gain & François, 2021). Non-significant parameters (*i*.*e*. degrees of freedom effectively shrunk to 0) were removed from the model. To test robustness of the parameters in the best fit model we used neighborhood cross validation with 45km radius. This radius was selected based on a correlogram of the residuals with 100km bins, taking the first distance that intersected with 0. We also looked for spatial autocorrelation through visual inspection of the residual correlogram and direct testing with Moran’s I test via the R package spdep (Pebesma & Bivand, 2023).

## Results

### Genome-wide ancestry is predictive of host preference

We first confirmed that a K of 3 is relevant for our dataset from NGSadmix results (Supp. Fig. 2) and that admixture between these major ancestry groups is prolific across the native range of *Ae. aegypti* (Supp. Fig. 3). We find that the proportion of human-specialist ancestry genome-wide is predictive of human host preference (Fig. 1B; Supp. Fig. 8, Supp. Fig. 9 R^2^ = 0.75), confirming predictions based on preliminary sampling (Rose et al., 2020). We find a non-linear (edf = 2.3; AIC = -54.67) relationship between host preference and human-specialist ancestry. This is more likely to be local plateauing than a global threshold as populations with a higher human-specialist ancestry than the maximum value in our models (e.g. invasive lineages in Orlando and Tahiti; (Rose et al., 2020) and populations from Rabai before 2014; (McBride et al., 2014)) do demonstrate even greater preference for human hosts.

**Figure 1.**
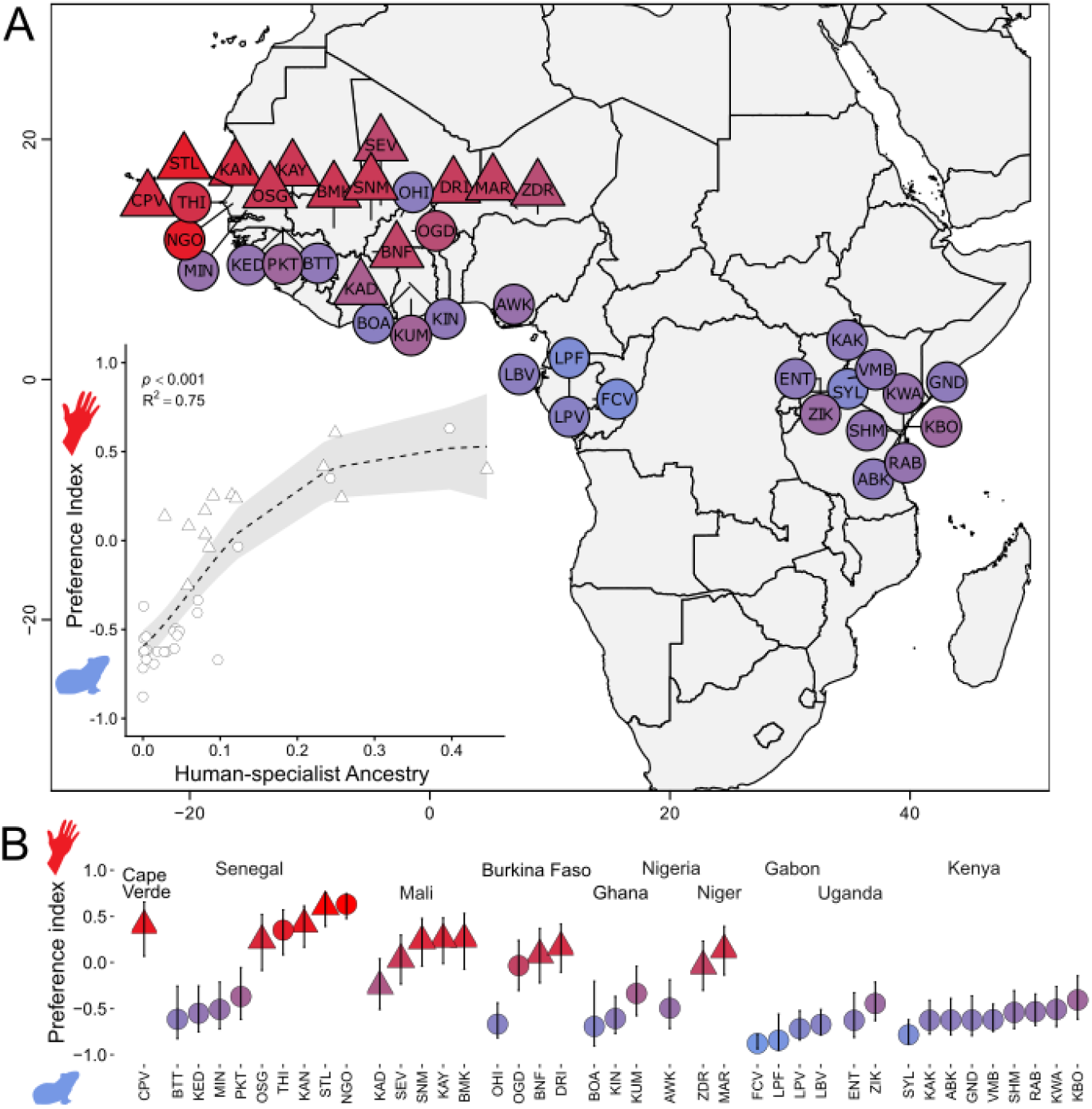
Human-specialist genome-wide ancestry predicts host preference. A) Map of all *Ae. aegypti* populations for which we have host preference behavioral data. Populations are colored based on the preference index, from -1 (animal preferring, blue) to 1 (human preferring, red). Inset shows relationship between ancestry and preference index with the GAM’s best fit line (dotted) and 95% confidence interval (grey box). B) Preference indices organized by country. Vertical line shows 95% confidence intervals. Circle: preference data from Rose 2020, Triangle: novel preference data.

### Selected regions of the genome that show higher predictability of host preference

We estimated human-specialist ancestry in 10mb windows with our f4-ratio traces (Supp. Fig. 4). After running separate GAMs on each 10mb window and then iterating across our different percentile thresholds, we found six windows, several of which overlap-leaving 3 distinct regions, that maximized our power to predict host preference (Supp. Fig. 5A) and were robust to LOOCV (Supp. Fig. 5B). Averaged together, these six windows showed a strong association (R^2^ = 0.86; AIC = -70.72) between human-specialist ancestry and host preference (Fig. 2A,D; Supp. Fig. 10). These regions were robust to a sensitivity analysis where we changed the size of our windowed analysis (Supp. Fig. 18). Further model selection looking at each continuous significant window individually produced a model with just three windows (windows starting at chromosome 1 279100000, chromosome 3 319820000 and 326440000) these three windows as separate terms resulted in a R^2^ of 0.921 (AIC = -77.12).

**Figure 2.**
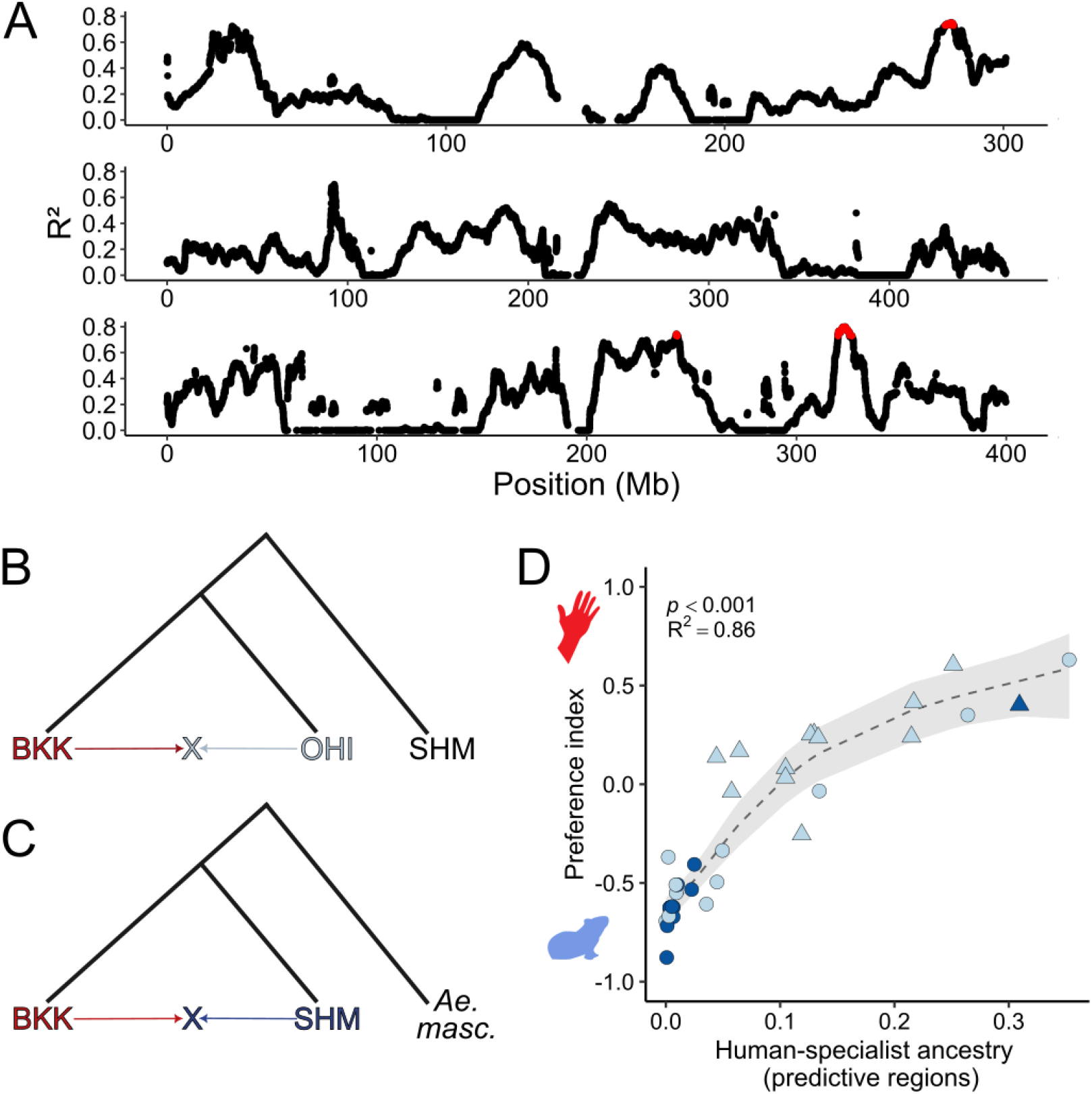
Human-specialist ancestry has outsized impact on host preference variation in selected regions of the genome. A) Manhattan plot showing the R^2^ from a GAM run at each individual window, red denotes windows that crossed the percentile (0.992) that maximized predictive power. B) F4-ratio tree topology for query populations (x) with generalist ancestry admixture from *AafW* only using equation: ((BKKa - SHM)*(X - OHI))/((BKKa - SHM)*(BKKb - OHI)) or C) admixture with *AafE* using the equation ((BKKa – Ae. masc)*(X - SHM))/((BKKa – Ae. masc)*(BKKb - SHM)). Human-specialist ancestry in red from BKK. D) Relationship between human-specialist ancestry from most predictive regions (*i*.*e*. red windows from A)) and host preference with the GAM’s best fit line (dotted) and 95% confidence interval (grey box). Light blue: admixture with *AafW*; dark blue: admixture with *AafE*.

**Figure 3.**
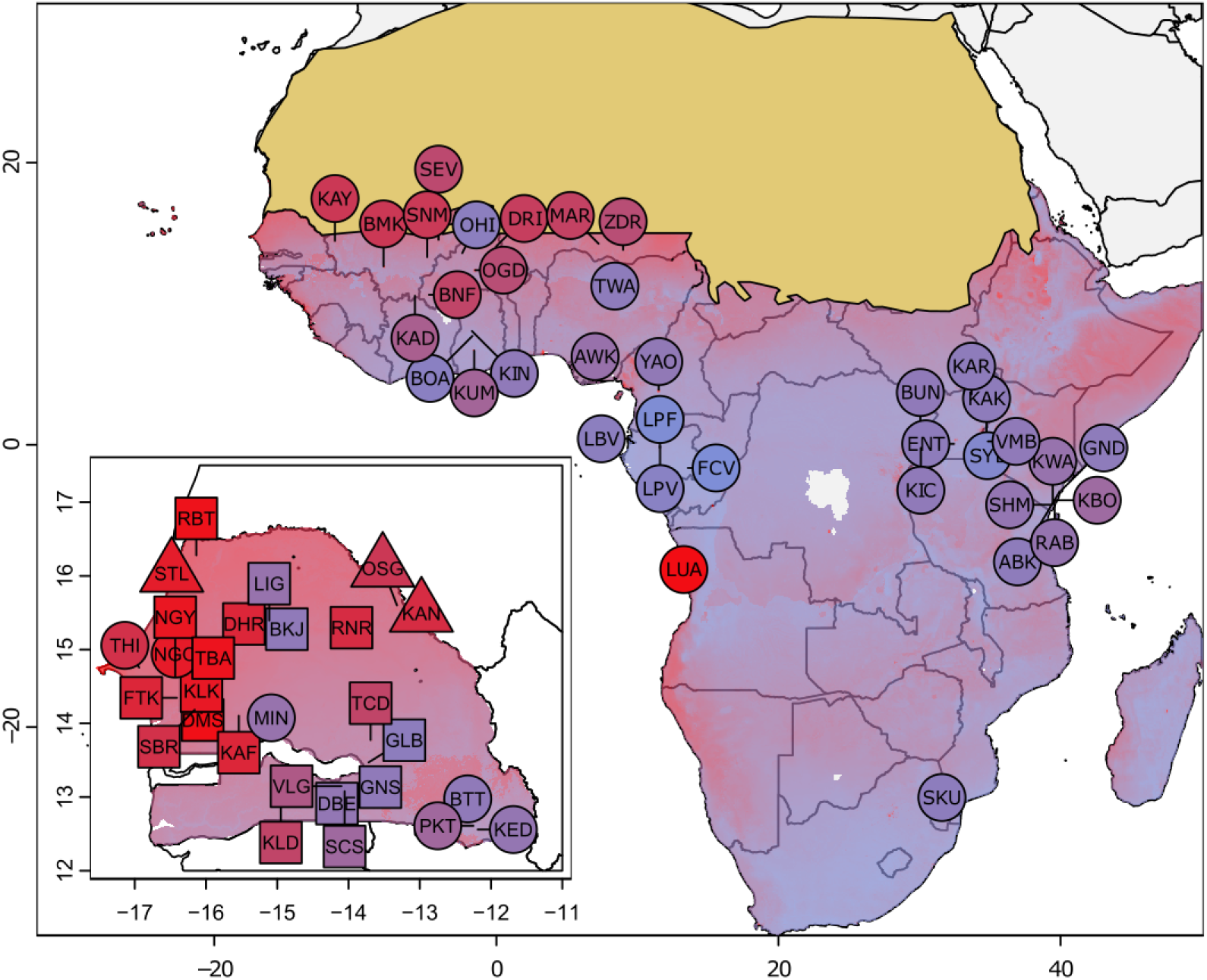
Predicted host preference (on the -1 to 1 preference index scale) based off our ecological model for populations with preference data (real, not predicted) containing the parameters bio9,15,17,19, npp, pet penman max, and dens 20. Inset shows Senegalese populations. Circle: preference data from Rose 2020; Triangle: novel preference data; Square: predicted preference from f4-ratio predictive windows. Tan overlay denotes Sahara desert-outside the contemporary range of *Ae. aegypti*.

Several potential genes of interest fall within the six predictive regions including genes known to be involved in host preference, most notability an odorant receptor (Or4) with well-established links to preference for humans (McBride et al., 2014), as well as ionotropic receptors IR93a and IR40a. IR93a is a nonolfactory IR coreceptor that drives human host proximity detection and blood-feeding behavior, Ir93a is required to seek high humidity associated with preferred egg-laying sites (Laursen et al., 2023). IR40a is a thermo- and hygrosensory neuron (THSN), specifically a “dry” THSN-meaning that it works in concert with Ir93a to detect dryness (Enjin et al., 2016). IR40a is also shown to be differentially expressed between human preferring and non-human preferring strains of *Ae. aegypti* (McBride et al., 2014). We also find several metabolic detoxification genes, known for their role in the metabolism of insecticides in insects (Li et al., 2007; Scott, 1999), including the carboxylesterase venom carboxylesterase-6-like gene and several cytochrome P450s including CYP304C1, CYP18A1, CYP306A1, CYP6AK1 and CYP6CD1.

While it is difficult to completely disentangle the causal effect of specific regions from the genome-wide signature, we attempted to do so by repeating the analysis but including whole genome ancestry as a covariate and also using LFMMs. Including genome-wide ancestry as a covariate does not add any new peaks and maintains the peak following 300mb on chr3 that we find when running this analysis without the genome-wide ancestry covariate (Supp. Fig. 14; Supp. Fig. 15). LFMM produced four significant loci (Supp. Fig. 6) which together still accounted for 76% of the variation in preference and did a better job of predicting preference than whole genome ancestry (Supp. Fig. 7). We looked for genes within 50kb upstream and downstream of the loci and found several-most notably one SNP was within a UTR region of the first exon of general odorant-binding protein 55 (OBP55). Several OBP genes within the OBP55 orthogroup have shown to be differentially expressed between *Culex pipiens pipiens* (avian preferring) and *Cx. pipiens molestus* (human preferring) populations (Noreuil & Fritz, 2021) and maxillary palps of *Anopheles quadriannulatus* (bovid preferring) and *An. coluzzi* (human preferring) (Athrey et al., 2017).

### Predicting preference from genomic information

We predicted preference for the samples for which we only had genomic information, using the six predictive windows from our f4-ratio analysis (Supp. Fig. 11). Several populations were predicted to have higher human host preference than any of the populations for which we have actual behavioral data (*i.e*. RBT, TBA, DMS, LUA and RABDOM). Of these, only one population, RABDOM, has any existing behavioral data (albeit with a different olfactometer setup) and our predictions (high human preference) agree with the measurements from that study (McBride et al., 2014). The predicted preference index for NGY (NGO but looking at larvae instead of adults) was 0.69, roughly similar to the actual preference for NGO adults at 0.63.

### Building ecological models predictive of host preference

When using GAMs to describe the variation in host preference, we find a model with bioclim2 variables mean temperature of driest quarter (bio9), precipitation seasonality (bio15), precipitation of driest quarter (bio17) and precipitation of coldest quarter (bio19), CHELSA variables net primary productivity (npp) and maximum potential evapotranspiration (pet penman max) and human density with a 20km radius buffer (dens 20) was the best fit for our preference data (R^2^ = 0.84, AIC = -64.36). All terms were approximately linear and the smooths for each term had a positive relationship with host preference with the exceptions of npp and pet penman max (Supp. Fig. 12). We find no signature of spatial autocorrelation both visually examining a correlogram of residuals at 100km bins and also using Moran’s I test (*p* = 0.39), but we still checked our best model with neighborhood cross validation (NCV), using neighborhood radius of 34km, selected by where the line passed 0 on the correlogram, and found our model was robust to NCV (R^2^ = 0.83, AIC = -58.27). Per chi-sq, bio15 seems to be the most important parameter in the model, and a simpler model with just bio15 and dens 20 and human density 20km radius buffer still does an adequate job of explaining the variance in preference, with bio15 switching from a linear to quadratic curve (R^2^ = 0.74, AIC = -54.59).

When expanding our preference dataset to include populations for which we have *predicted* preference from the genomic data, we find a model with bio15, maximum vapor pressure deficit (vpd max), and dens 20 best fit our data (R^2^ = 0.80, AIC = -86.42) with both bio15 and dens 20 as approximately linear and vpd max as a 4^th^ order polynomial. When going back and using this model to fit the actual preference dataset, we see this model performs worse than our full model (i.e. bioclim2 variables 9,15,17,19, CHELSA variables npp and pet penman max and dens 20), but about the same as the bio15 and dens 20 only model (R^2^ = 0.74, AIC = -54.59). When we use our full model from the actual preference only data to fit the expanded preference dataset, we find the full model still does an adequate job of explaining the overall variation (R^2^ = 0.74). However, the main difference between the full model and the model with bio15, vpd max and dens 20 is in Senegal where the full model does significantly worse (R^2^ = 0.57) compared to the bio15, vpd max and dens 20 model (R^2^ = 0.78) and this outperformance holds true even when only looking at Senegal populations with actual preference data (R^2^ = 0.82 versus R^2^ = 0.83) (Supp. Fig. 13; Supp. Fig. 17).

## Discussion

Here we demonstrate the feasibility of an ecological genomics approach to predict the distribution of a trait with significance to the rising disease burden of arboviruses. We show that 86% of the variation in host preference can be explained by ancestry at just six 10mb windows, and these windows contain both previously identified (e.g. Or4) and novel candidate genes (IR93a and IR40) that could be influencing host preference. Frequencies at four loci also do a respectable job (75%) of predicting host preference and reveals a potential host preference candidate gene in OBP55. However, it should not be lost that whole-genome ancestry only performs slightly worse than the four loci. This seems to be indicative of linkage among genetic variation that dictates a wide range of behaviors important for urban adaptations. This is consistent with the observed divergence between the two ecotypes in many traits relevant for thriving in urban environments (reviewed in (Fifer et al., 2025)) including differences in egg temperature tolerance (Chakraborty et al., 2024) to deal with higher urban temperatures, and adult oviposition preferences (Peterson, 1977; Xia et al., 2021) and greater hatching success (Metz et al., 2023) in artificial container environments.

We demonstrate the feasibility of building a predictive ecological model in this system, with major contributions from precipitation seasonality and human density in explaining how host preference is partitioned across Africa. Other less important terms in the model seem to explain the variance in a subset of the populations, for example precipitation of the coldest quarter appears to predict variation in host preference for populations outside of the Sahel region. We also show that increasing our behavioral data sample sizes by including the populations for which we make genotype-based-preference predictions and then re-running our model selection, maintains the importance of precipitation seasonality and human density, but also provides a model with a more robust prediction in Senegal (the country whose sample size increased the most with the addition of these genotype-based-preference populations) through the inclusion of vapor pressure deficit. When including the vapor pressure deficit term, we can more clearly see the divide between human preferring and more generalist populations in Senegal which seems to occur over central Senegal when environments transition to more forested areas.

The Sahel region presents an interesting area of investigation for the origin of human specialization behavior in *Ae. aegypti*. Sahelian populations exhibit both the greatest variation in host preference between populations and include the most human preferring populations in our sampled range. Sahel’s high precipitation seasonality, driven by its long, up to 9-month dry seasons, presents a challenge for the mosquito to complete its water-obligate life cycle, but artificial sources of water buffer against this seasonal limitation. This dynamic is consistent with our findings that both human density and precipitation seasonality heavily contribute to explaining the distribution of human preferring populations contemporarily, but additionally offers possible insights into the origin of human preference. The Sahelian climate has been largely consistent for the last 5,000 years following the end of the African humid period and the drying of the Sahara (Tierney et al., 2017). This environmental transition also coincides with the divergence time between generalist and human-specialist ancestry, supporting a Sahelian origin of human specialization (Rose et al., 2023). However, explicit testing between a Sahelian origin and other alternate origin hypotheses (e.g. Angolan; (Powell et al., 2018)) is still an important area for future research.

Together our ecological genomic approach to understanding the distribution of host preference in *Ae. aegypti* offers several key insights. We demonstrate the feasibility of predicting the preference of mosquito populations from genotype alone, substantially reducing the phenotyping burden associated with generating data on a public-health-relevant trait. We also show that this data-collection obstacle can be reduced even further, by predicting the host preference of populations with only environmental data. This framework provides a guide for addressing data gaps in disease vectors and offers information that can be used to identify regions at risk under current and future environmental conditions.

## Acknowledgements

This work was supported as part of a Global Collaborative Network through the Princeton Institute of International and Regional Studies, NIH K22 AI166268 to NHR, and a Helen Hay Whitney Foundation Fellowship to JEF. The authors thank Lindy McBride for her support and advice about this work.

## Supplemental Material

**Supplemental Table 1.**
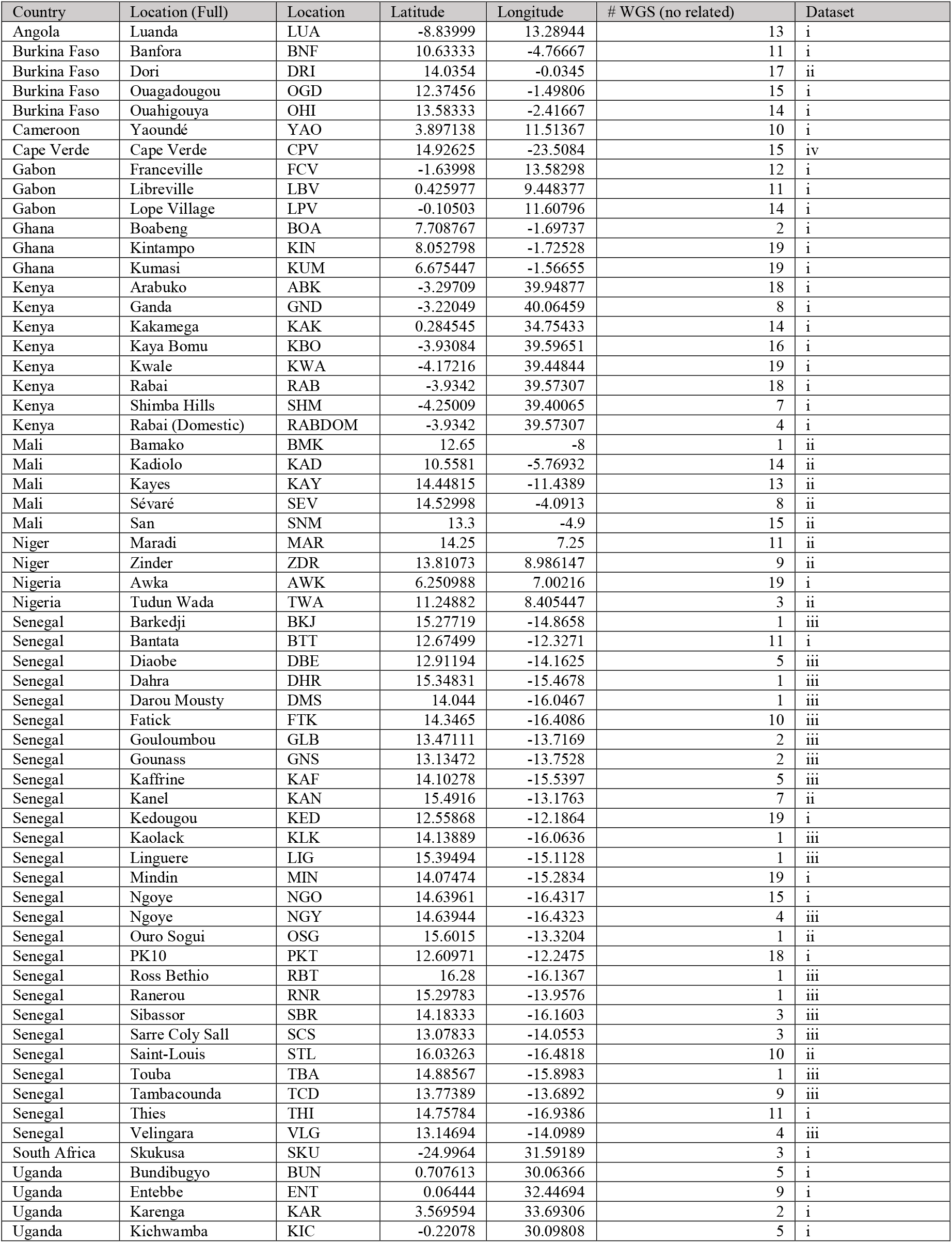
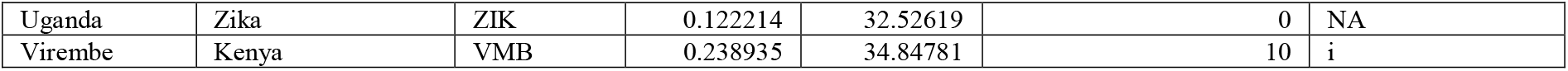
All populations used in this study. Dataset key: (i) ∼10-15x coverage WGS dataset of adult female and male *Ae. aegypti* from the 1206 *Ae. aegypti* genomes project (Crawford et al 2025). (ii) 1-2x coverage WGS of adult female and male Aedes aegypti from 2021 collections. (iii) 1-2x coverage WGS of larval Aedes aegypti from 2020 rainy season. (iv) ∼10-15x coverage WGS dataset of adult female and male *Ae. aegypti* from Cape Verde (Rose et al 2022).

**Supplemental Figure 1.**
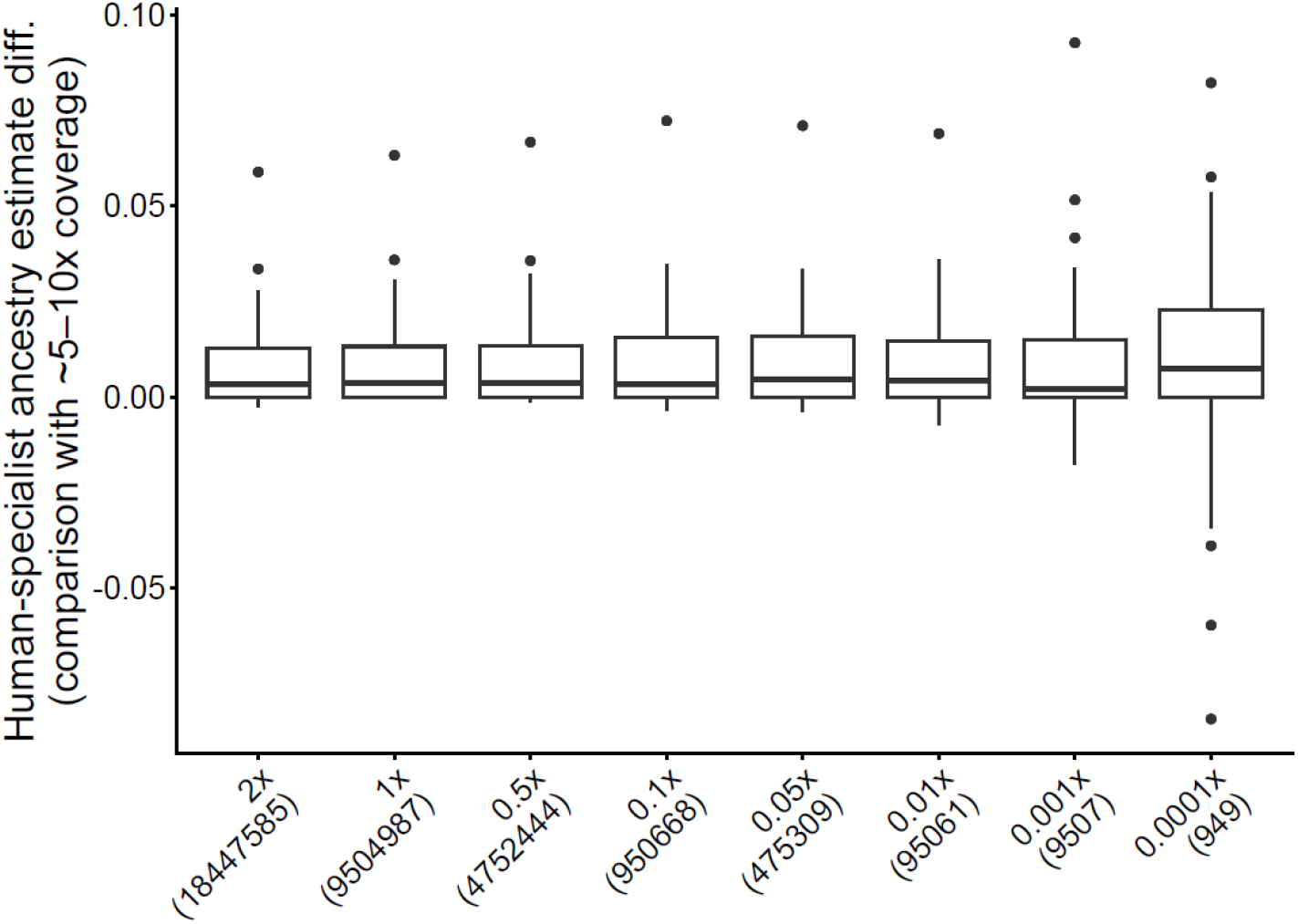
FastNGSadmix estimates of human-specialist ancestry are robust to very low coverage data. Box plots show estimate difference in comparison with higher coverage (i.e. 5-10x) samples. Boxes show the interquartile range (25th–75th percentiles) with the median indicated by the central line; whiskers extend to the most extreme non-outlier values, and points represent outliers. Coverage calculated in reference to chromosomal reference with zeros included. Average chromosomal reads as shown in parentheses.

**Supplemental Figure 2.**
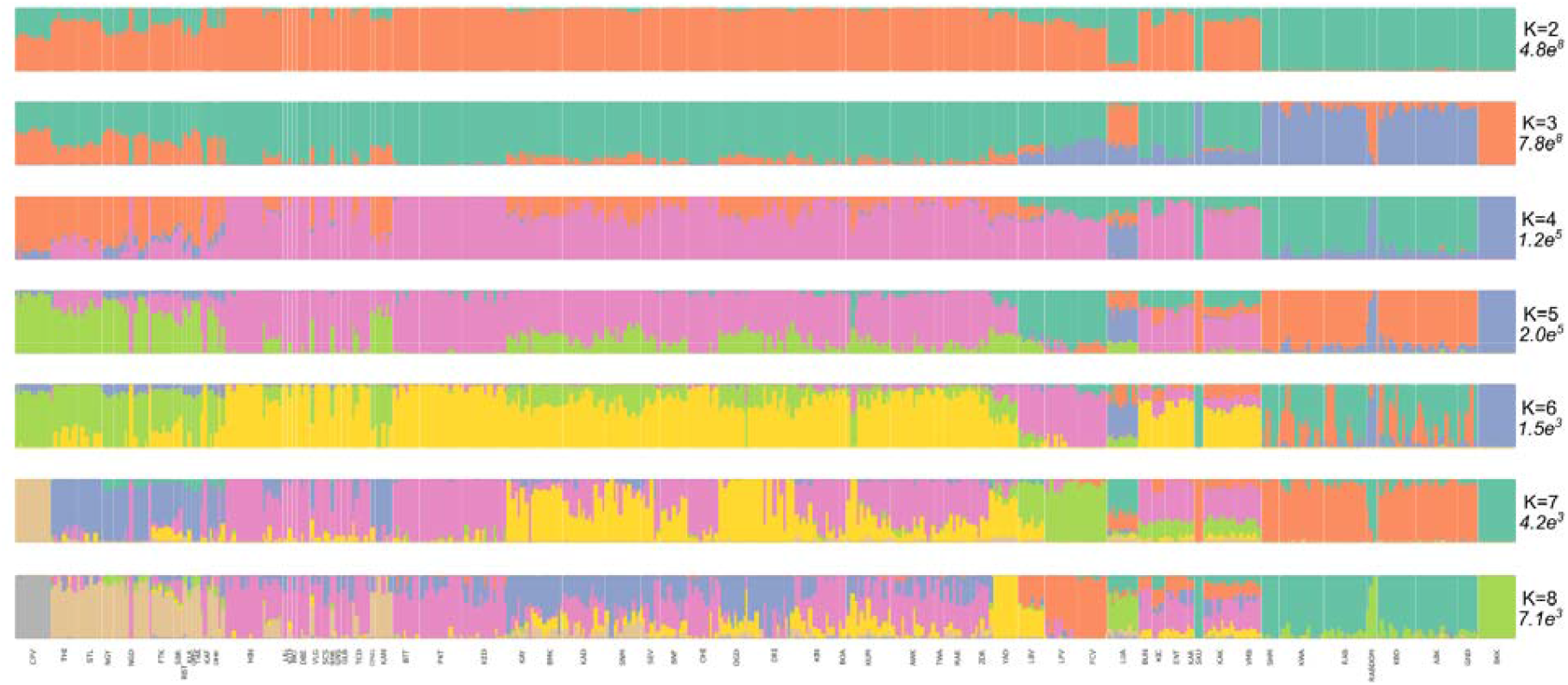
NGSadmix estimates for all populations showing K of 2:8; each K’s likelihood is shown in italics.

**Supplemental Figure 3.**
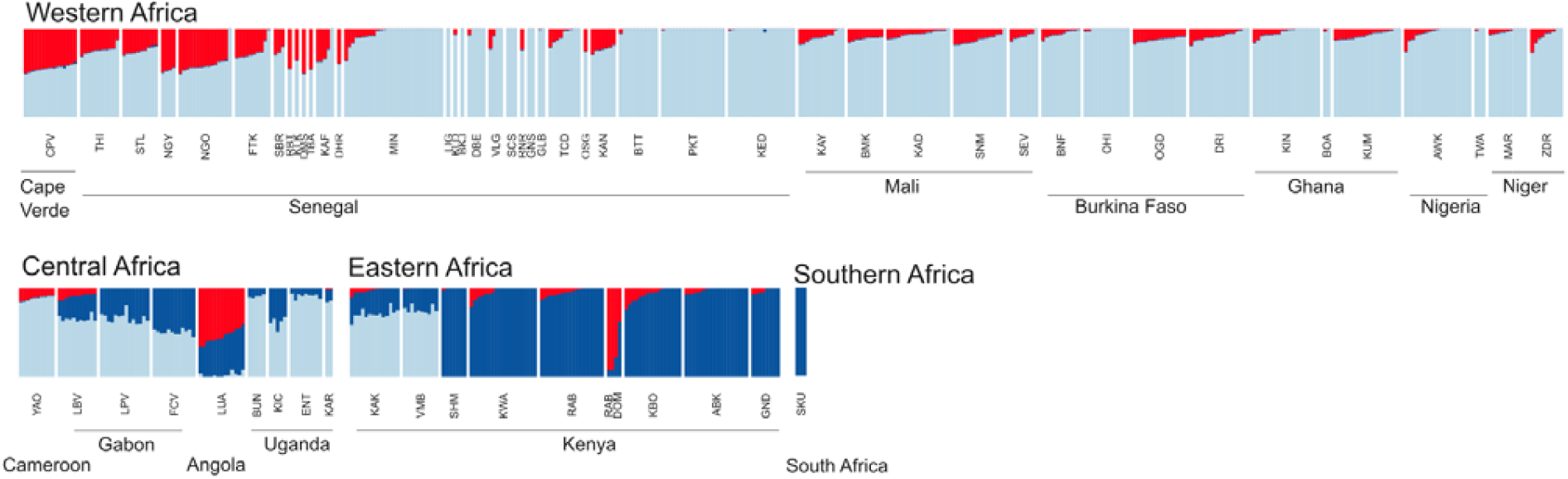
FastNGSadmix estimates for all populations using the K of 3 reference panel. Red: human-specialist ancestry; light blue: AafW ancestry; dark blue: AafE ancestry.

**Supplemental Figure 4A.**
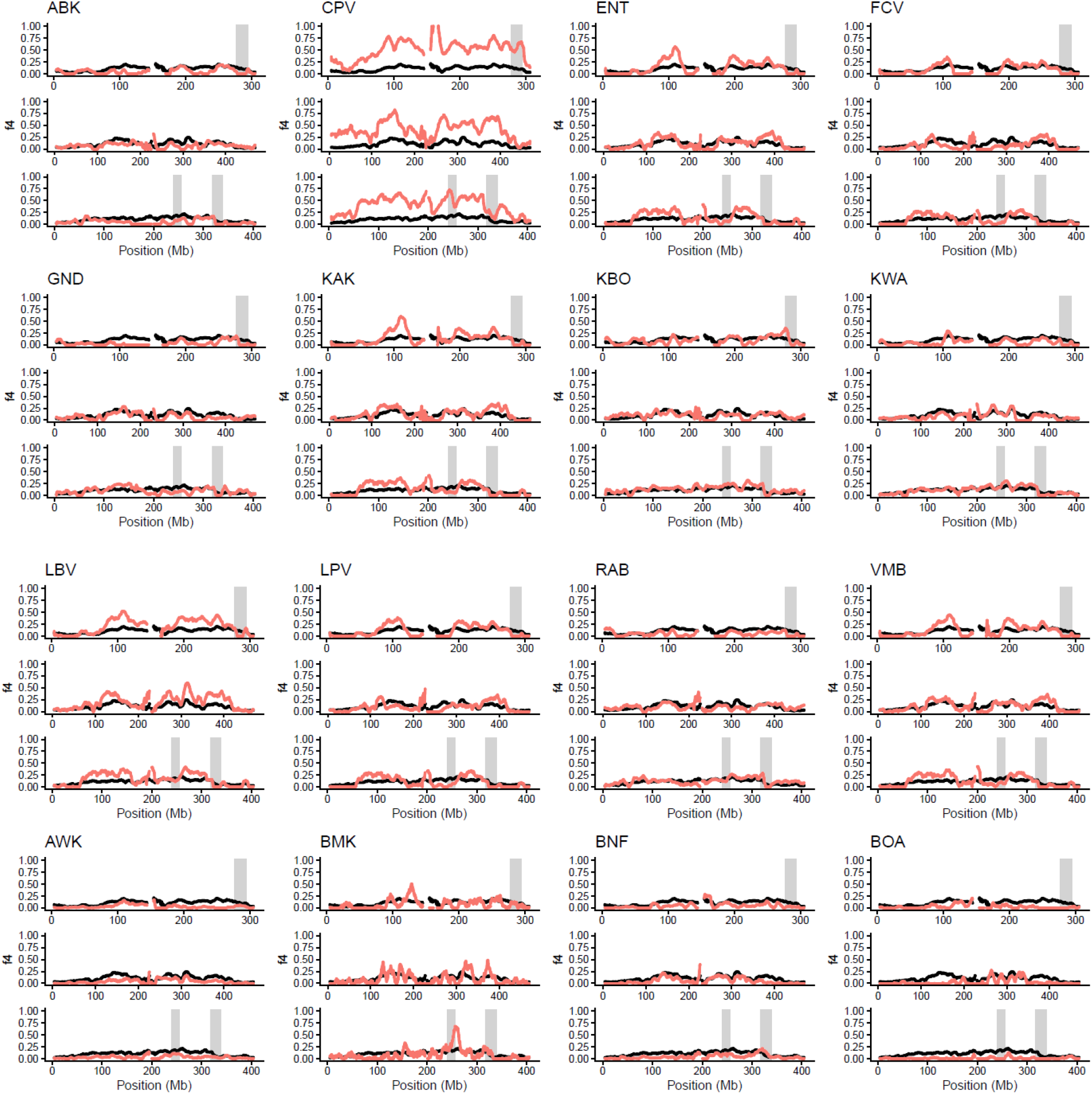
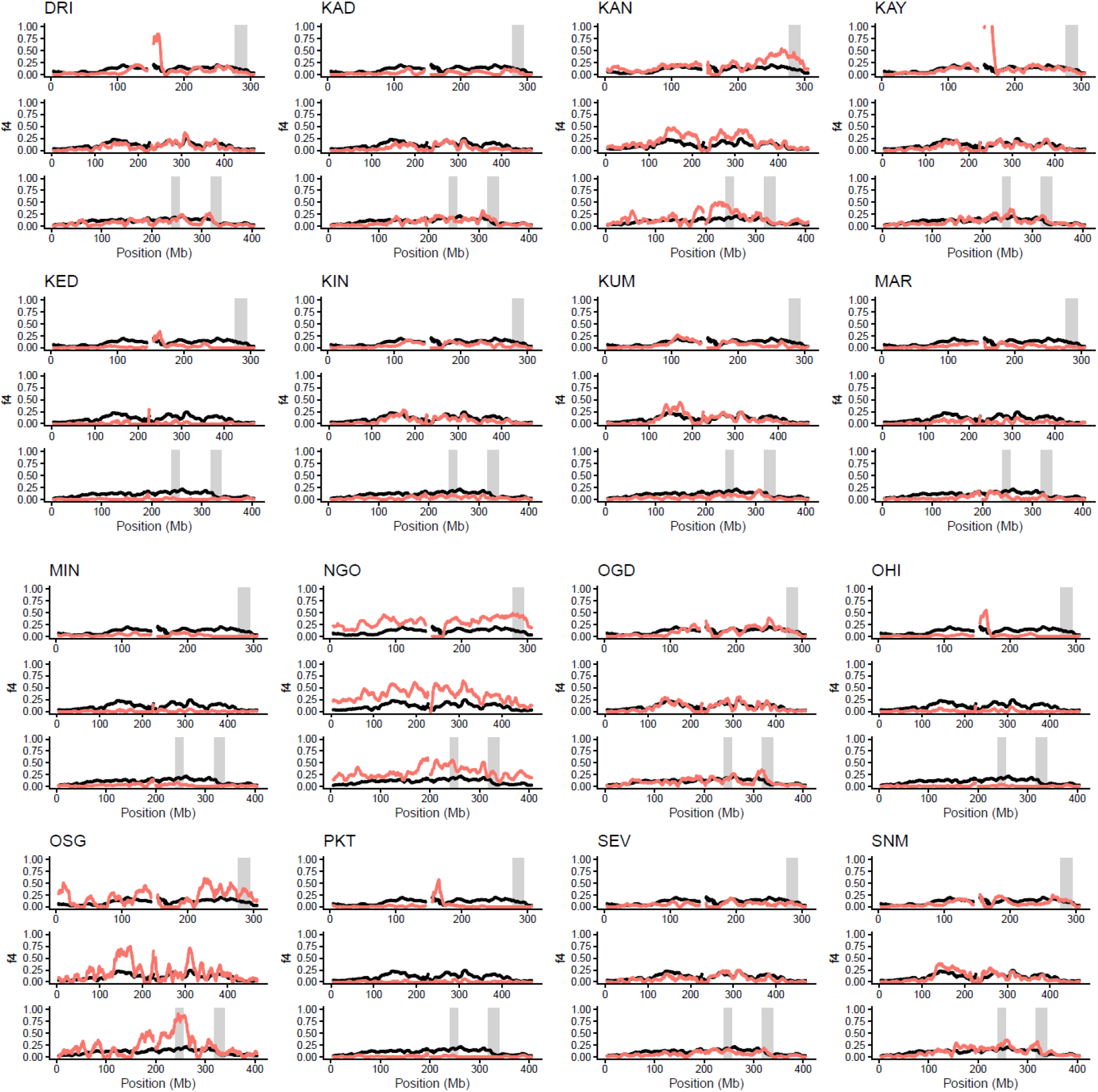
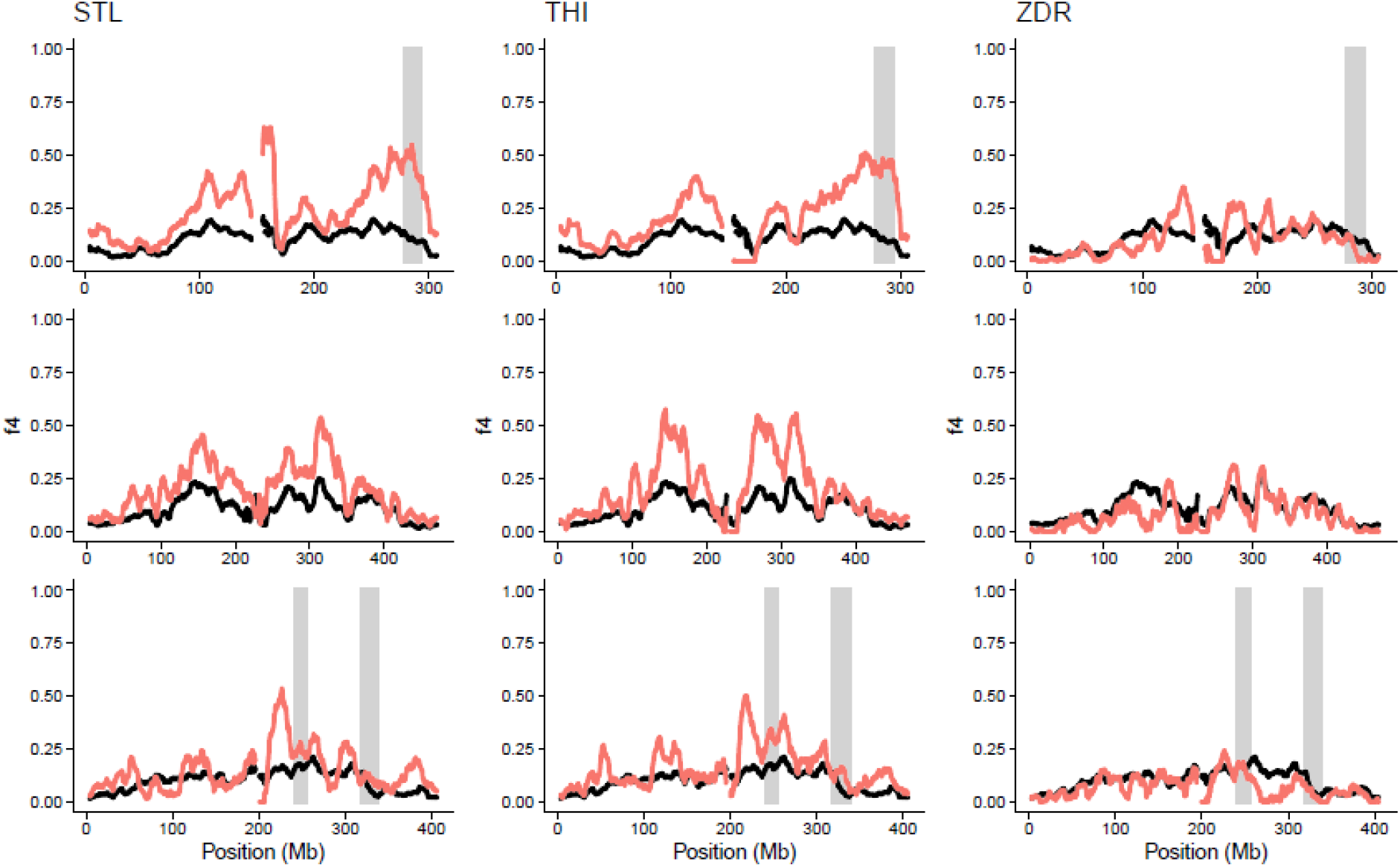
Human-specialist ancestry trace calculated in 10mb windows using f4-ratio estimates. Red line: f4-ratio traces for each query population with population behavior; black line: averaged f4-ratio for all populations with preference data. Grey boxes: most predictive regions (*i.e*. red windows from Fig. 2A). Top to bottom: chromosomes 1,2,3.

**Supplemental Figure 4B.**
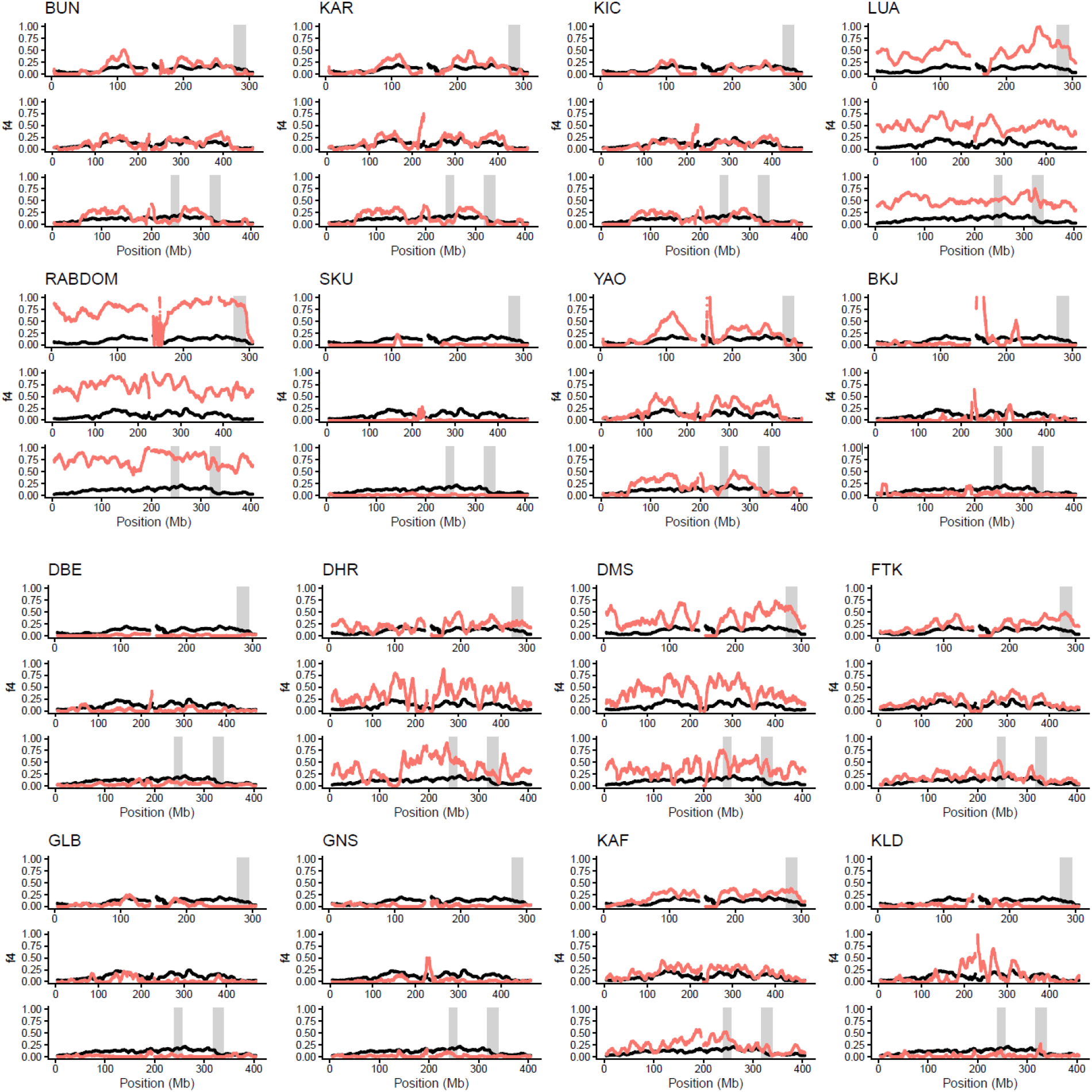
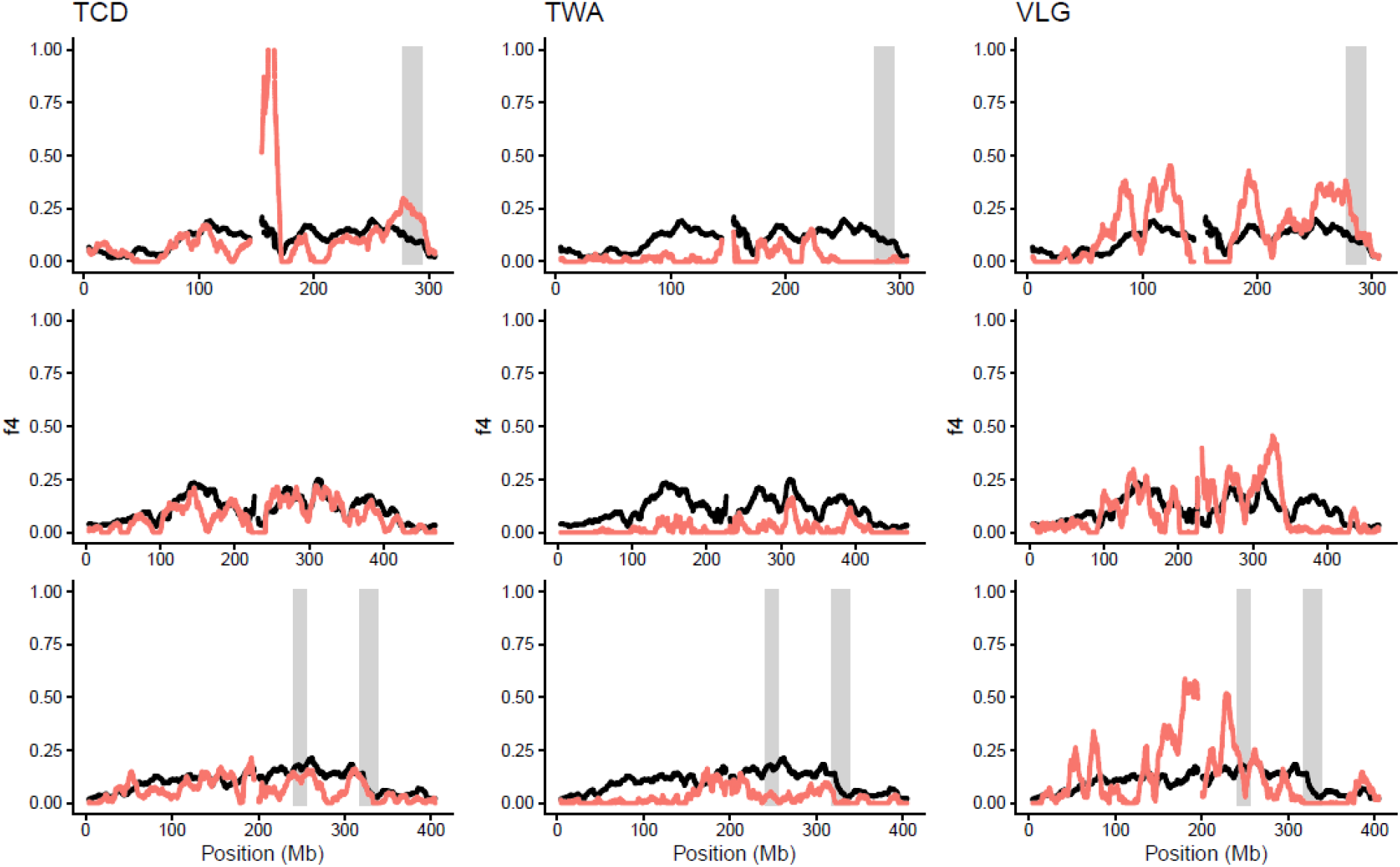
Human-specialist ancestry trace calculated in 10mb windows using f4-ratio estimates. Red line: f4-ratio traces for each query population without population behavior; black line: averaged f4-ratio for all populations with preference data. Grey boxes: most predictive regions (*i.e*. red windows from Fig. 2A). Top to bottom: chromosomes 1,2,3.

**Supplemental Figure 5.**
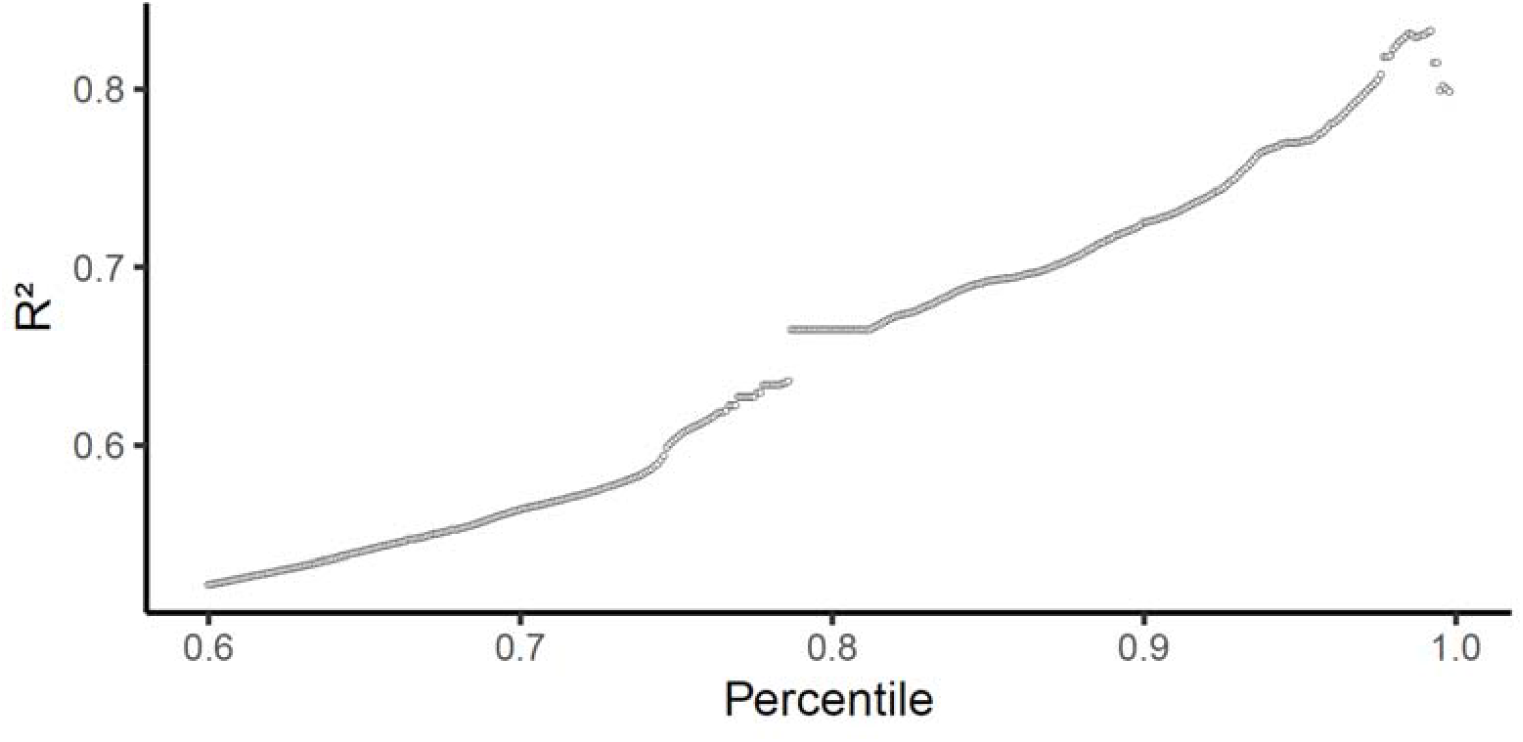
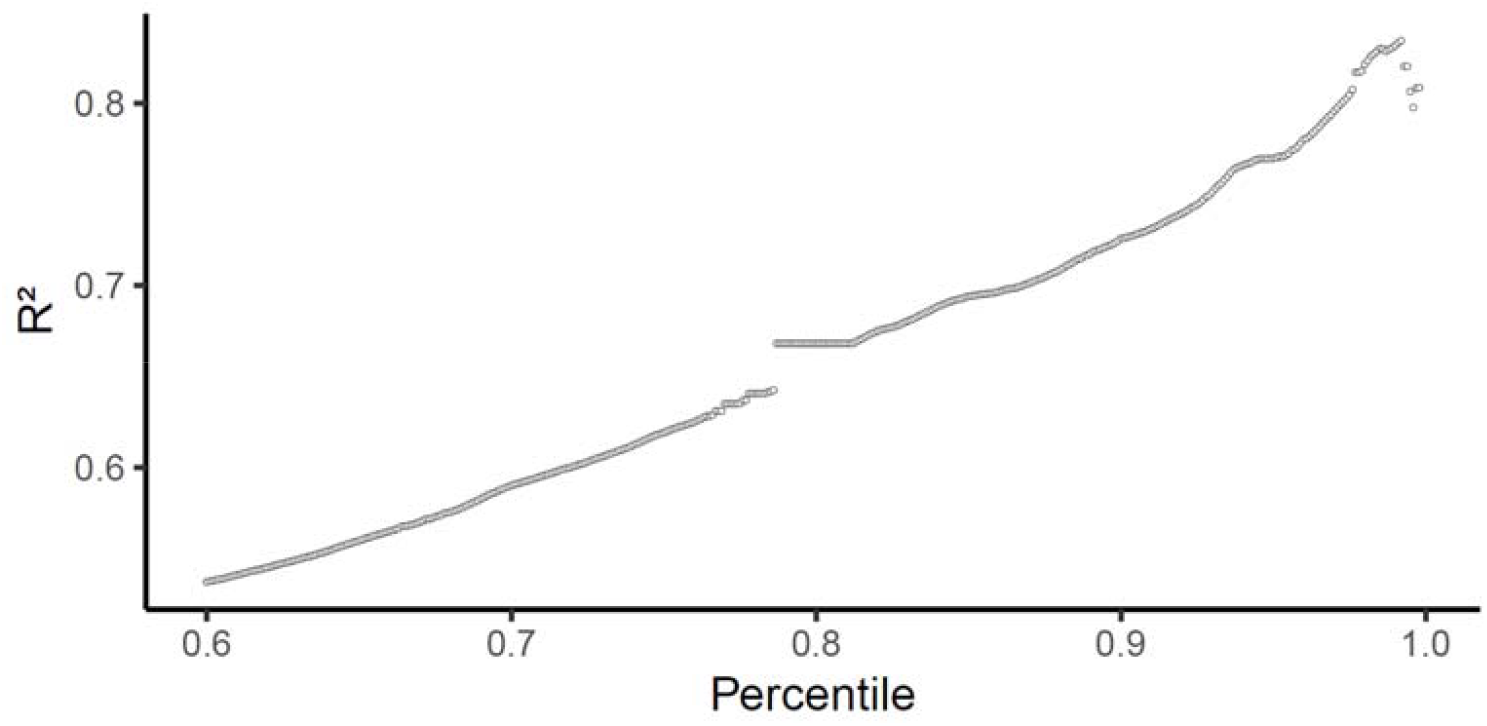
Finding the genomic regions that maximize predictive power of human-specialist ancestry for human host preference. Top) Plot shows predictive power, using a Gaussian GAM on logit transformed preference data of an entire genomic region when 10mb windows are collapsed according to different R^2^ percentile thresholds (0.60-0.999, increasing by 0.001). Bottom) Analysis repeated using a LOOCV cross validation.

**Supplemental Figure 6.**
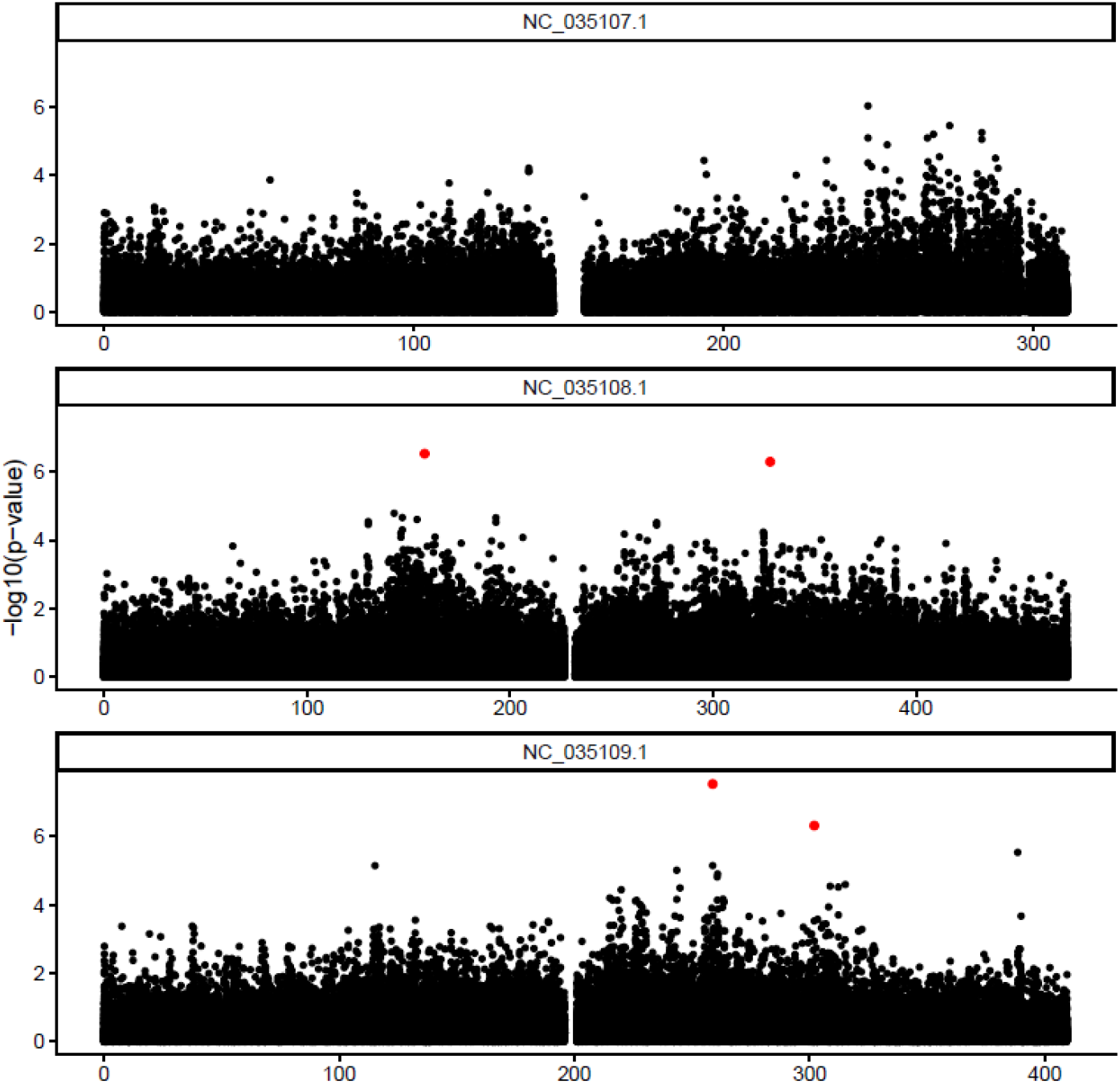
Manhattan plot showing LFMM results identifying loci associated with human host preference. P-values were calibrated using genomic control and adjusted for multiple testing using FDR, with a significance threshold of 0.05 (red). P-values are shown on a −log10 scale.

**Supplemental Figure 7.**
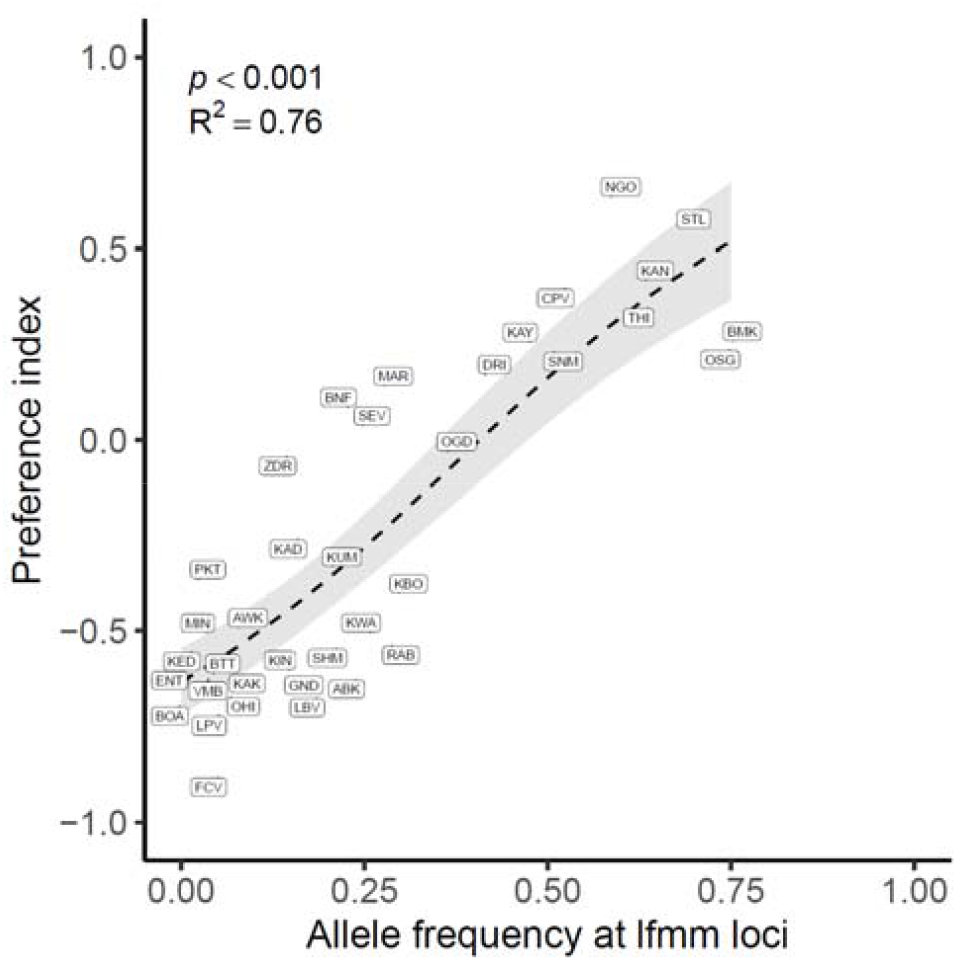
Allele frequencies at the four significant LFMM loci (Supp. Fig. 6) predict human host preference. Dotted line shows GAM model fit, grey denotes approximate 95% confidence intervals calculated as the predicted value ± 1.96 × standard error.

**Supplemental Figure 8.**
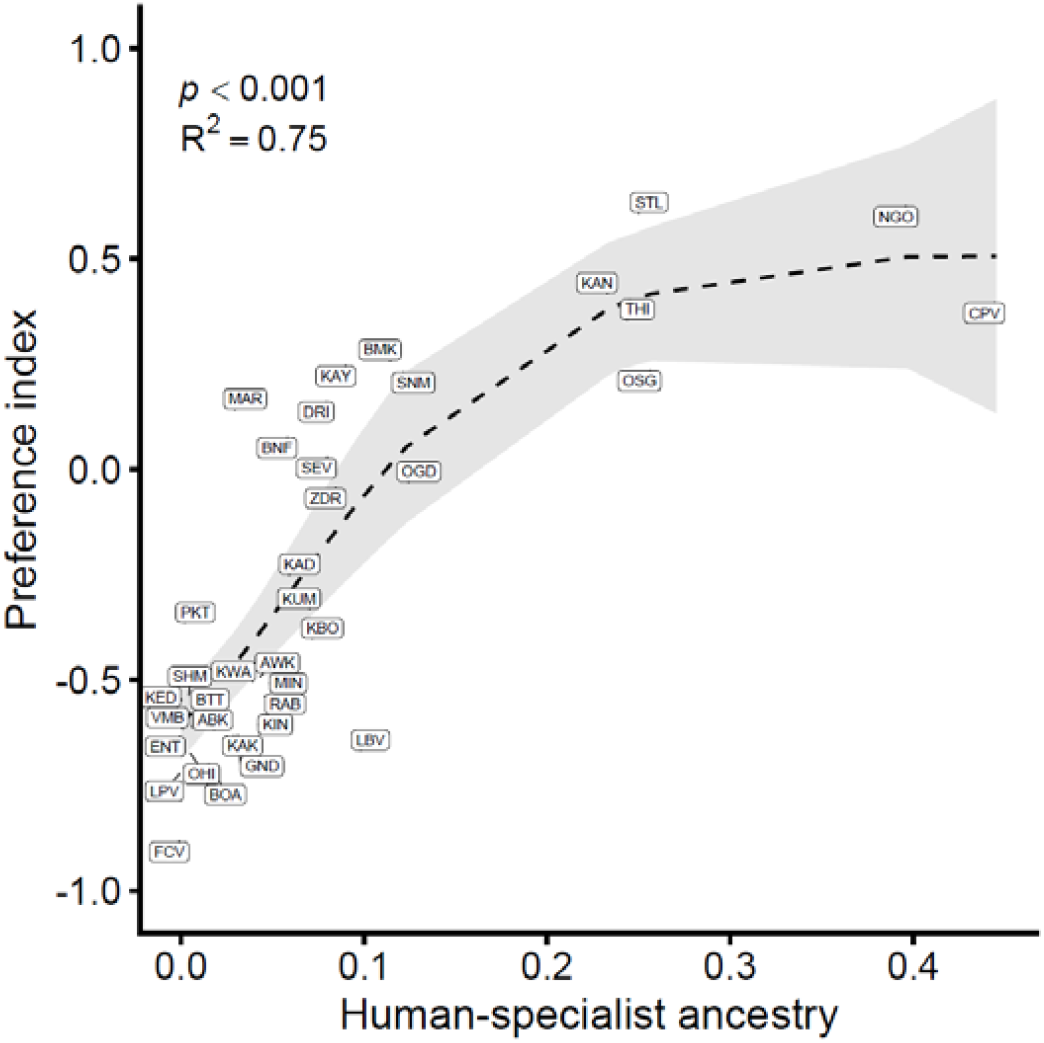
Genome-wide ancestry predicts human host preference. Plot is the same as Fig. 1A, but shows population labels. Dotted line shows GAM model fit, grey denotes approximate 95% confidence intervals calculated as the predicted value ± 1.96 × standard error.

**Supplemental Figure 9.**
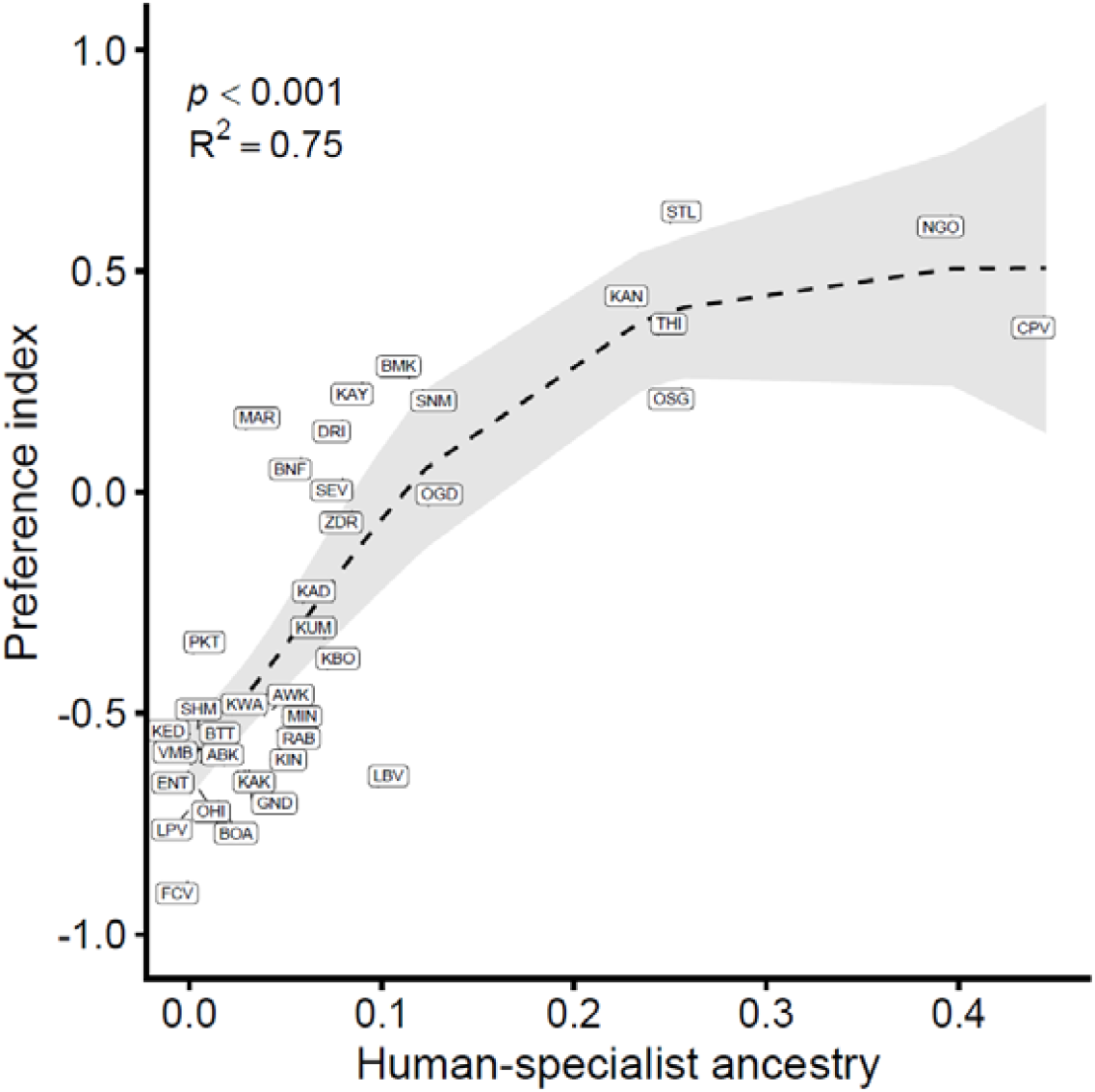
Genome-wide ancestry human host preference predictions are robust to LOOCV. R^2^ and p-value shown are derived from LOOCV. Dotted line shows GAM model fit, grey denotes approximate 95% confidence intervals calculated as the predicted value ± 1.96 × standard error.

**Supplemental Figure 10.**
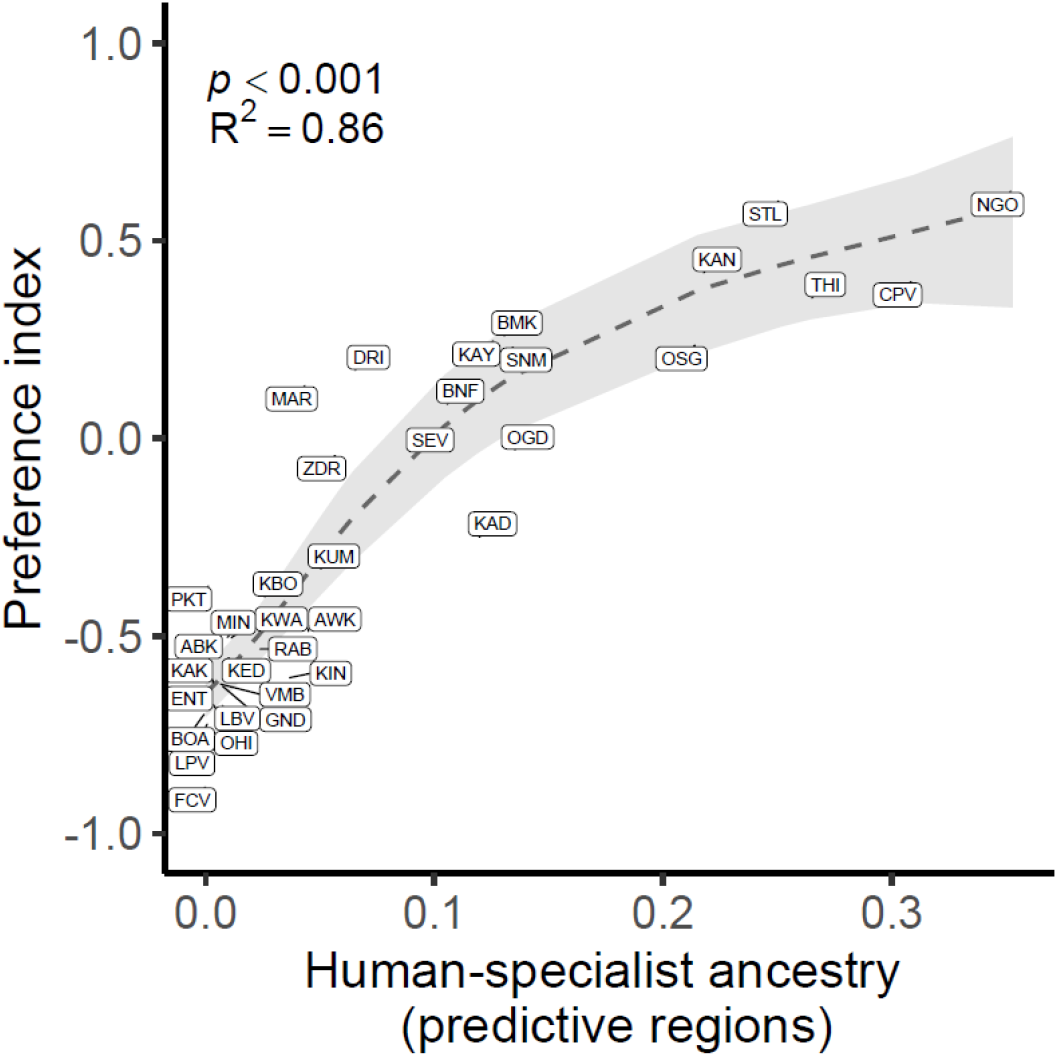
Relationship between human-specialist ancestry from the most predictive regions and host preference with the GAM’s best fit line (dotted) and 95% confidence interval (grey box). Plot is modified version of Fig. 2B, replacing circles with population labels.

**Supplemental Figure 11.**
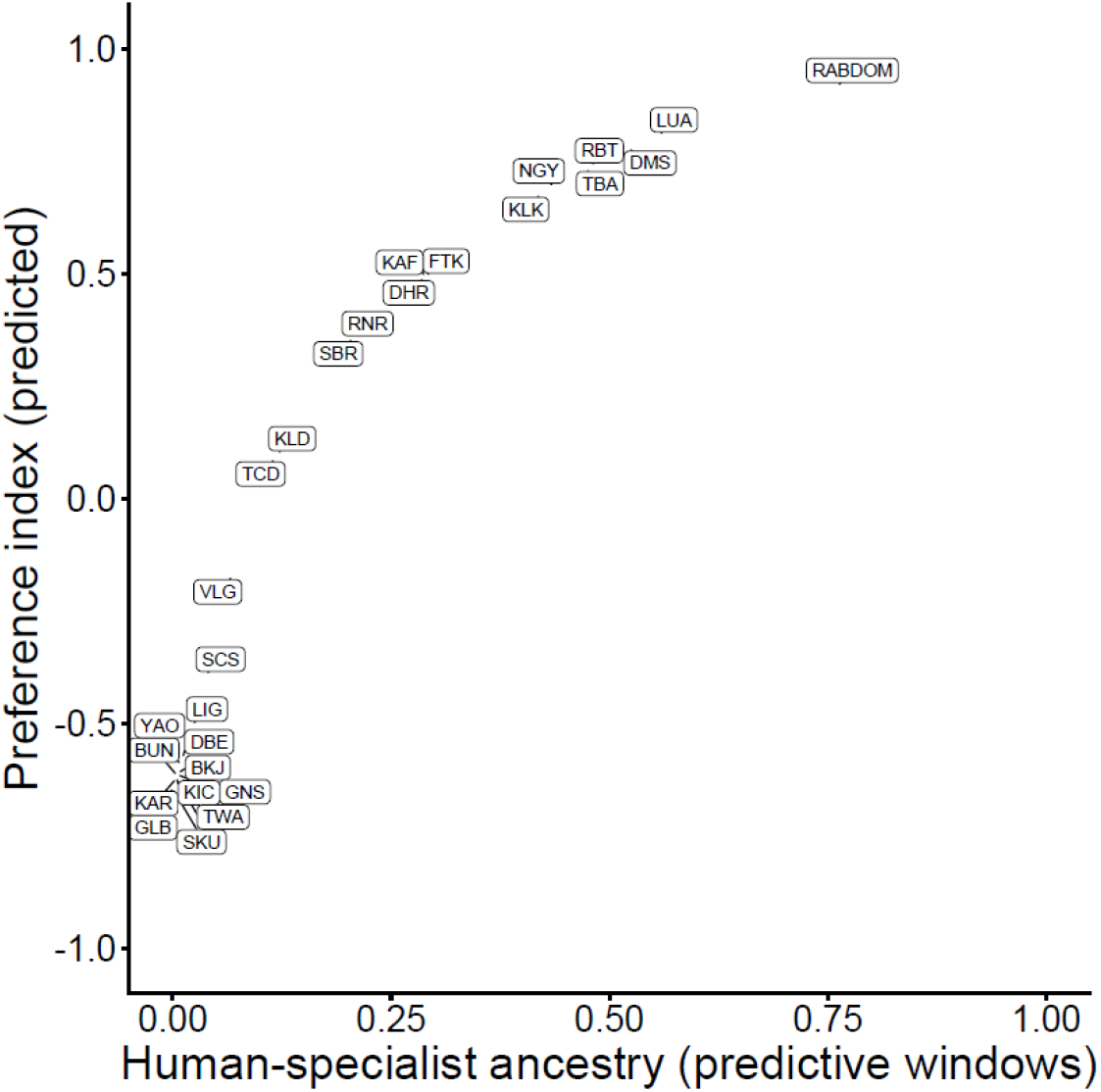
Genomic based host preference predictions. Predictions are made using the predictive regions from Fig. 2. Plot shows all populations for which there is genomic data, but no actual preference data.

**Supplemental Figure 12.**
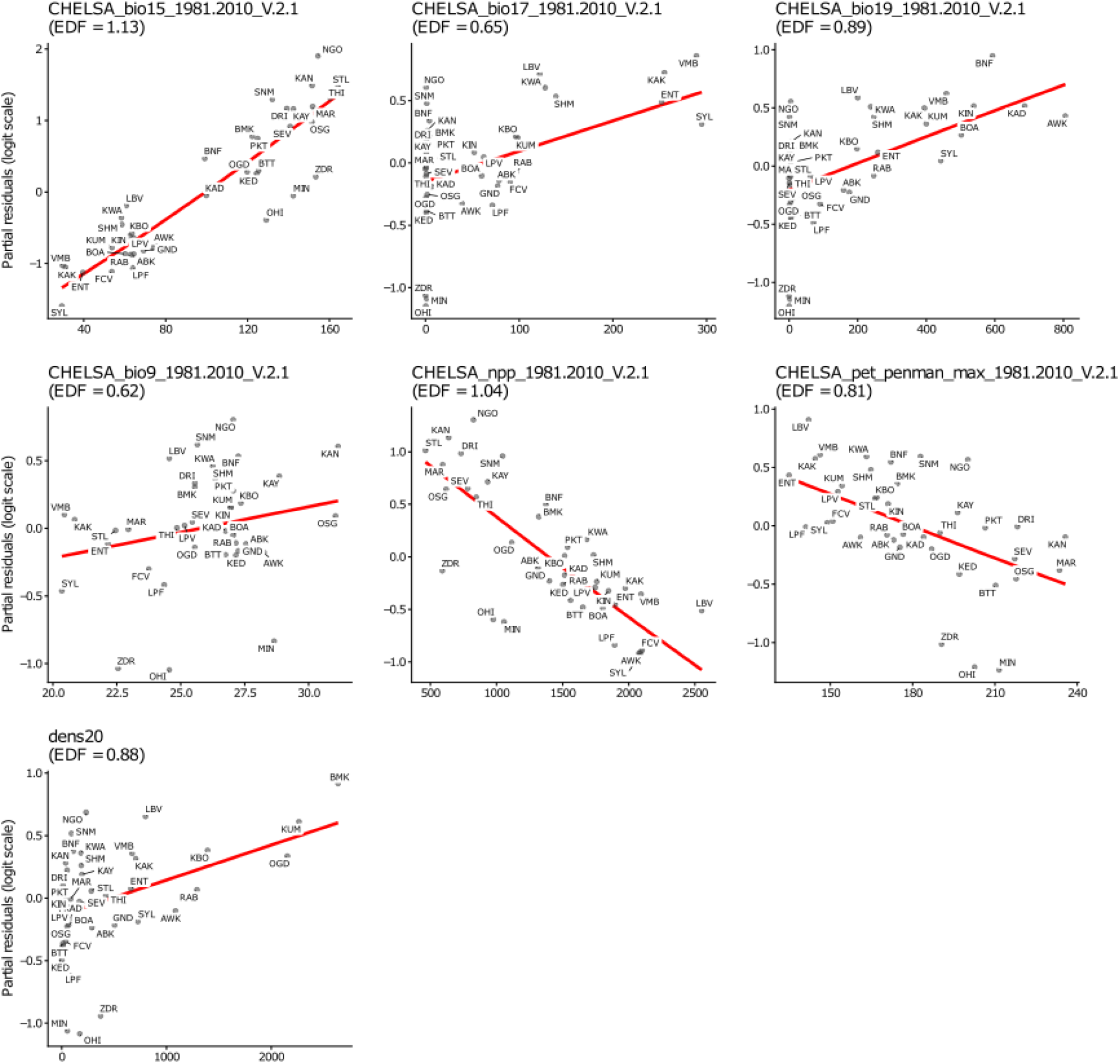
Breaking down individual significant terms from the ecological model predicting host preference (Fig 3). Red line: partial effect fit; dots: partial residuals for each population.

**Supplemental Figure 13.**
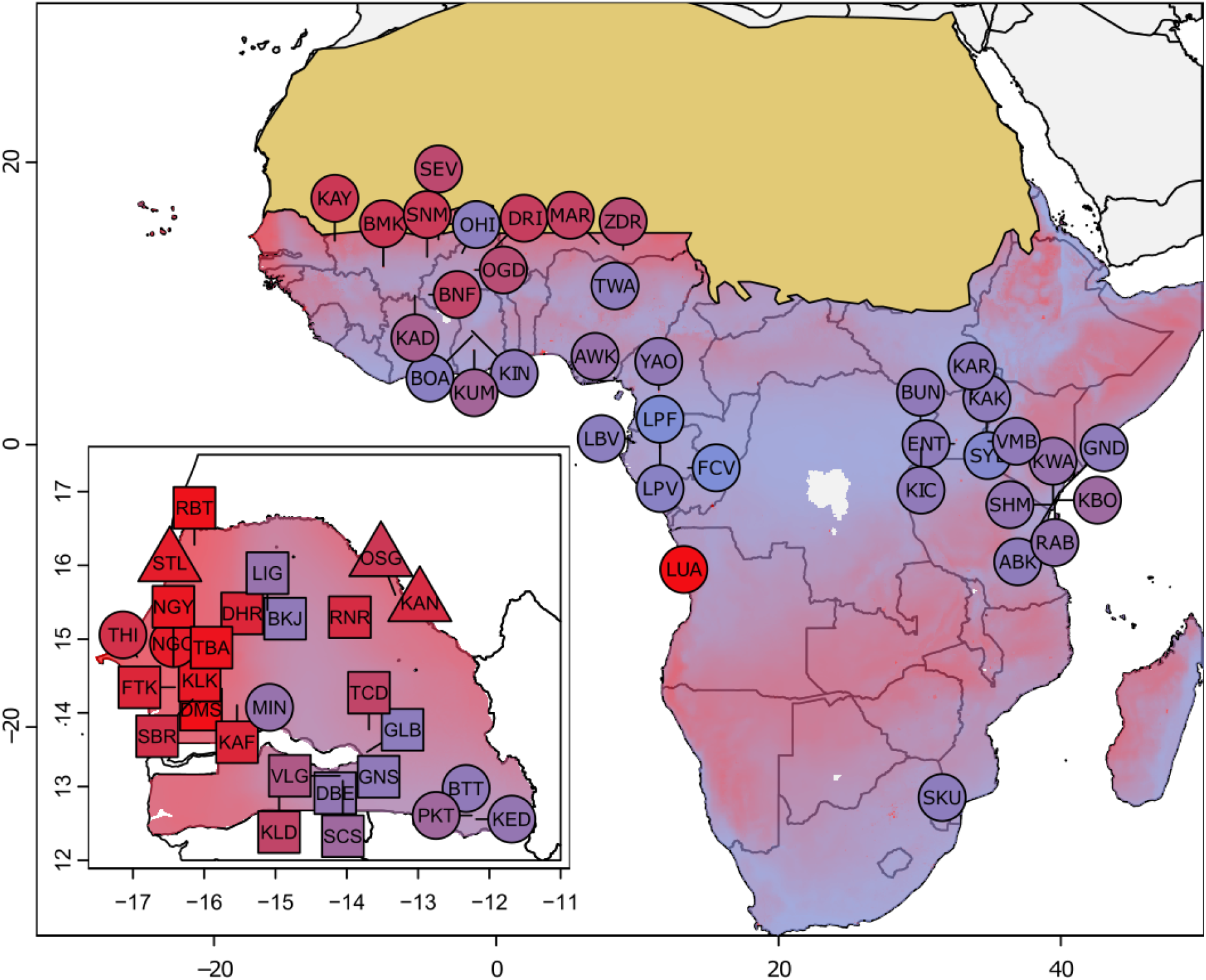
Predicted host preference (on the -1 to 1 preference index scale) based off our ecological model on all populations (actual and genotype predicted preference) containing the parameters bio15, max vpd, and dens 20. Inset shows Senegalese populations. Circle: preference data from Rose 2020; Triangle: novel preference data; Square: predicted preference. Tan overlay denotes Sahara desert-outside the contemporary range of *Ae. aegypti*.

**Supplemental Figure 14.**
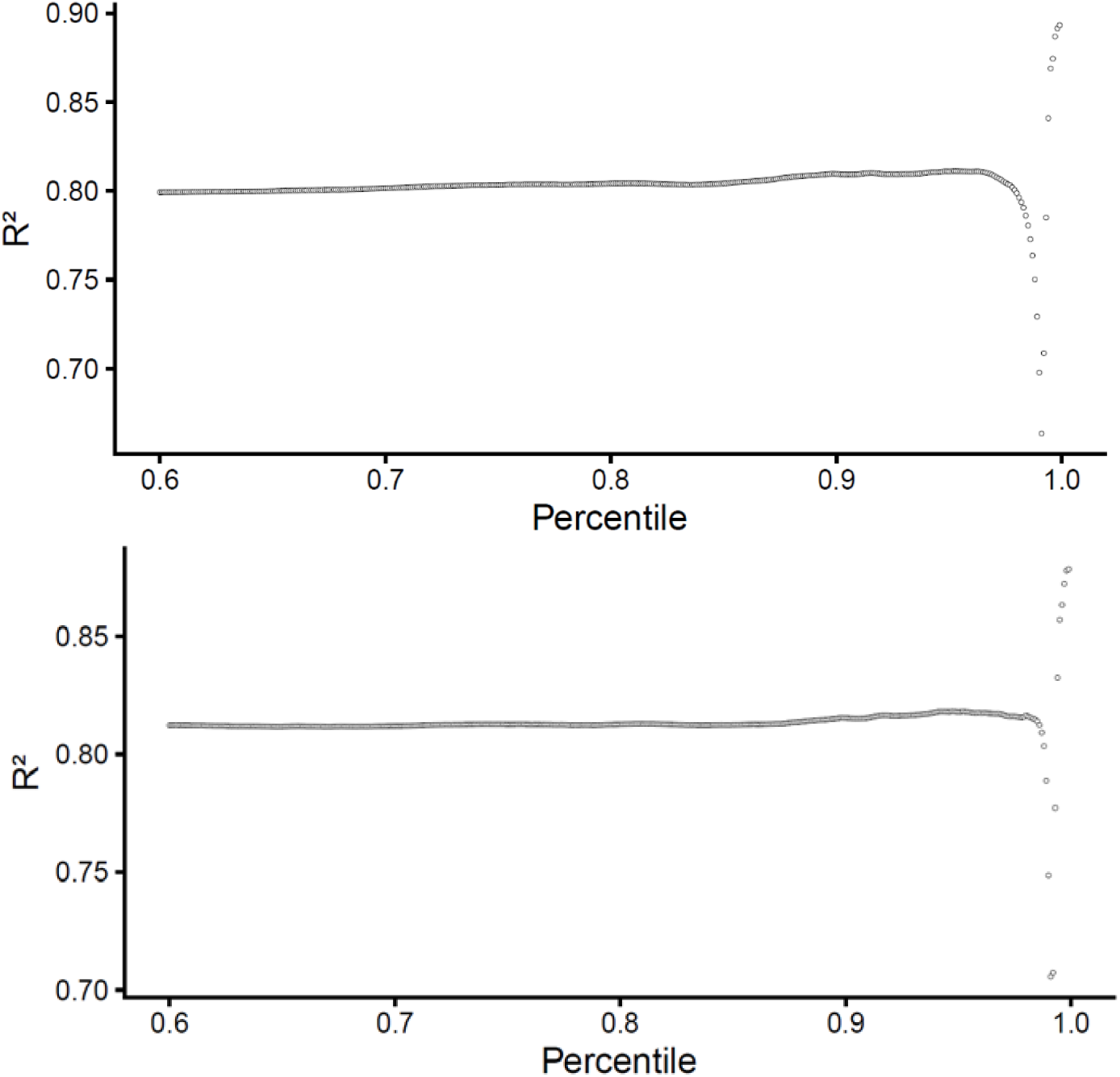
Finding the genomic regions that maximize predictive power of human-specialist ancestry for human host preference while controlling for whole-genome ancestry, by including it as a covariate in the GAM. Top) Plot shows predictive power, using a Gaussian GAM on logit transformed preference data of an entire genomic region when 10mb windows are collapsed according to different R^2^ percentile thresholds (0.60-0.999, increasing by 0.001). Bottom) Analysis repeated using a LOOCV cross validation.

**Supplemental Figure 15.**
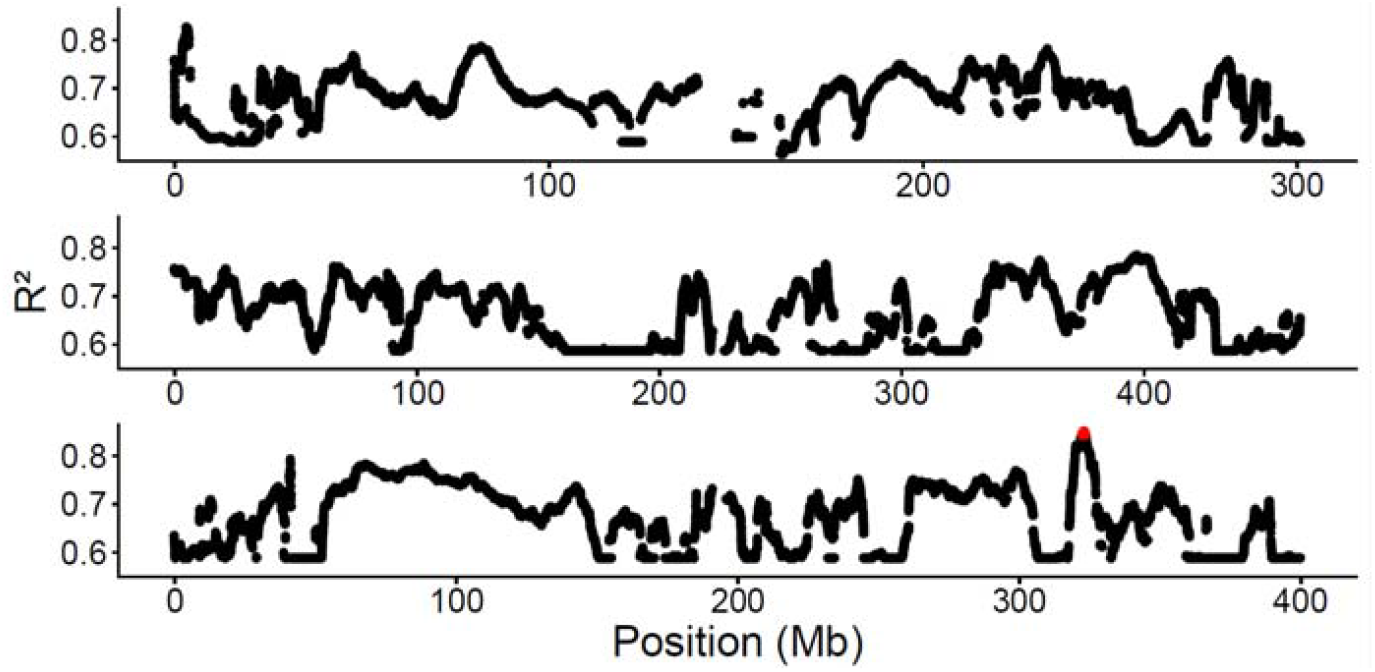
Finding the genomic regions that maximize predictive power of human-specialist ancestry for human host preference while controlling for whole-genome ancestry by including it as a covariate in the GAM. Manhattan plot showing in red the most predictive R^2^ percentile from Supp. Fig. 14 (0.999).

**Supplemental Figure 16.**
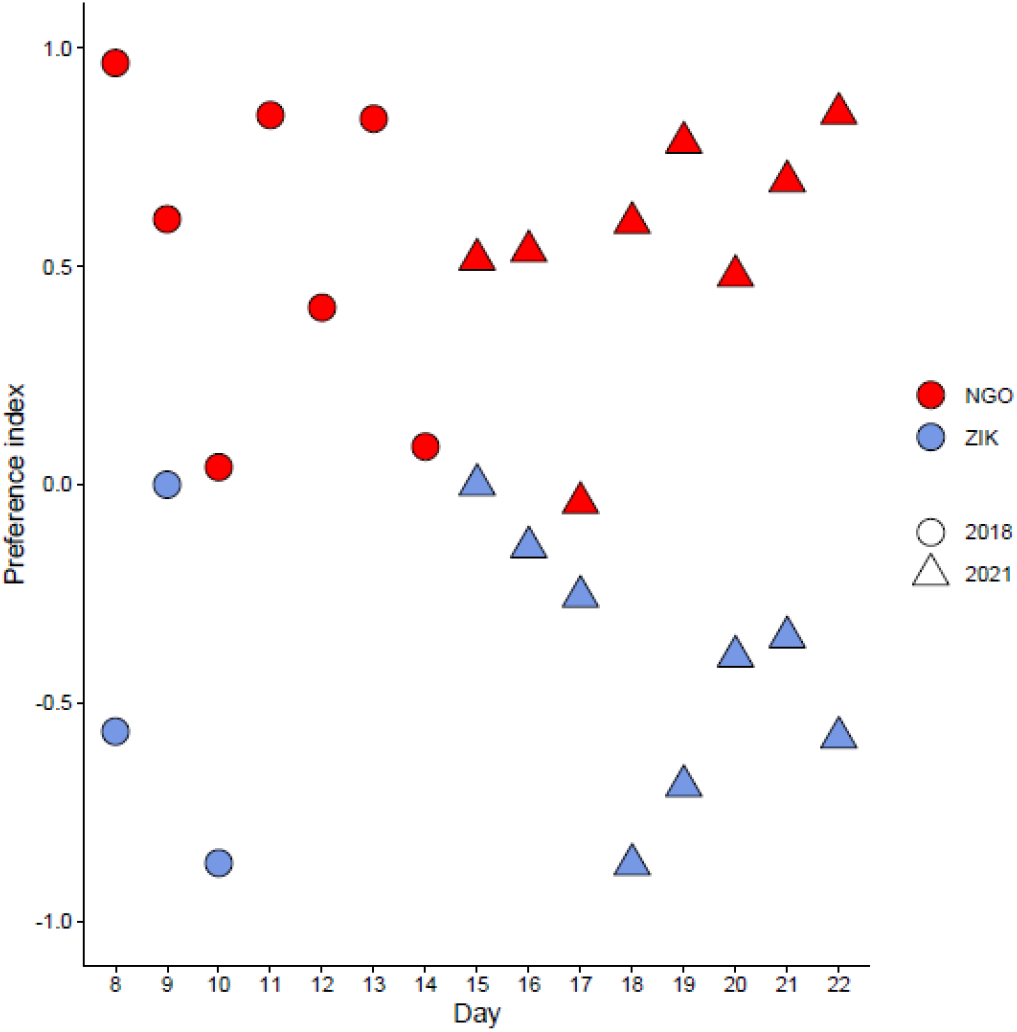
Behavioral data for colonies used across the 2018 and 2021 behavioral experiments, plotted by daily preference.

**Supplemental Figure 17.**
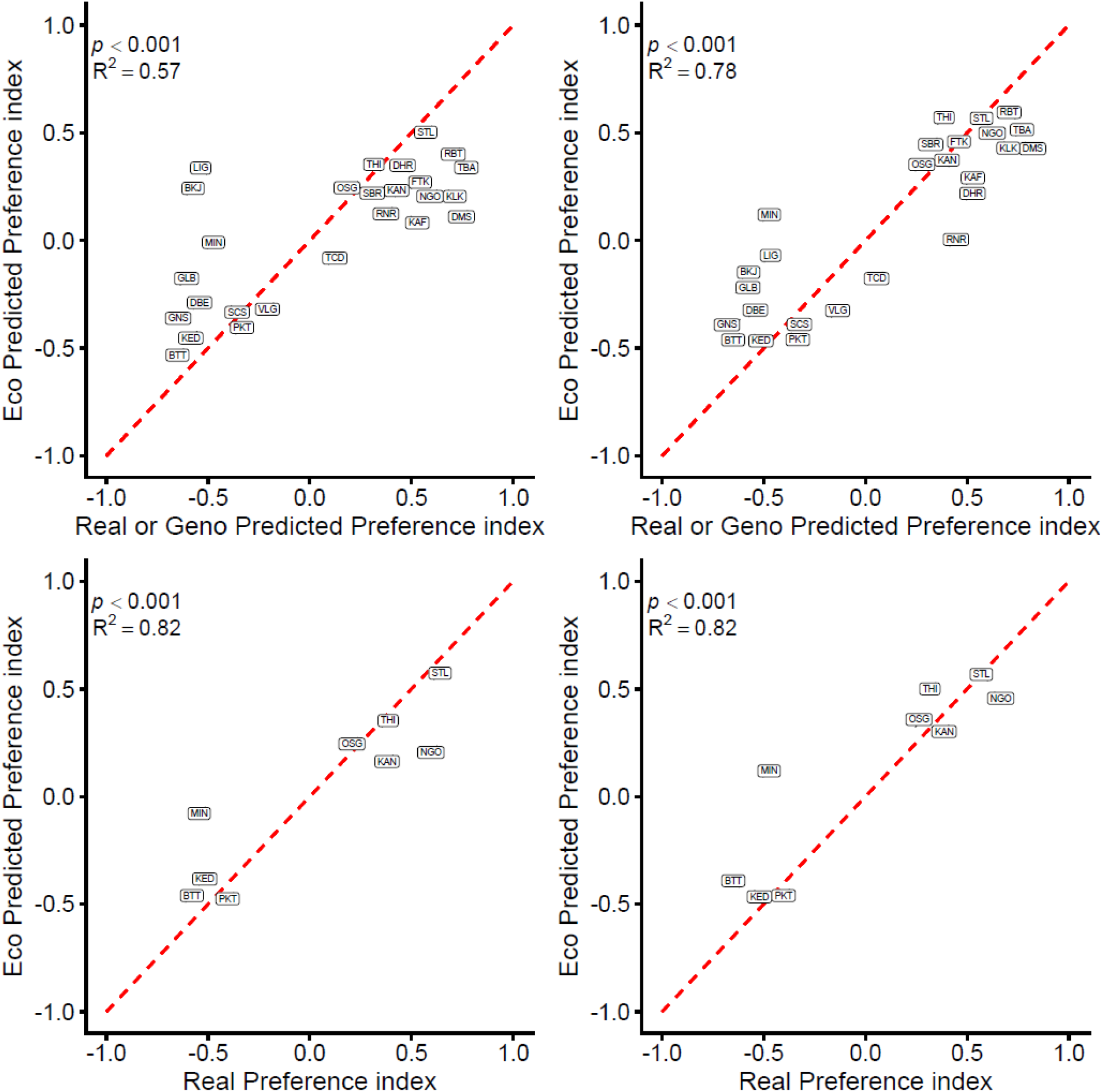
Comparison between the two ecological models for predicting preference in Senegal. Top: All Senegalese populations; Bottom: Only Senegalese populations that have real preference data; Left: Real preference model (bio9,15,17,19, npp, pet penman max, and dens 20) ; Right: Expanded preference model (bio15, vpd max and dens 20).

**Supplemental Figure 18.**
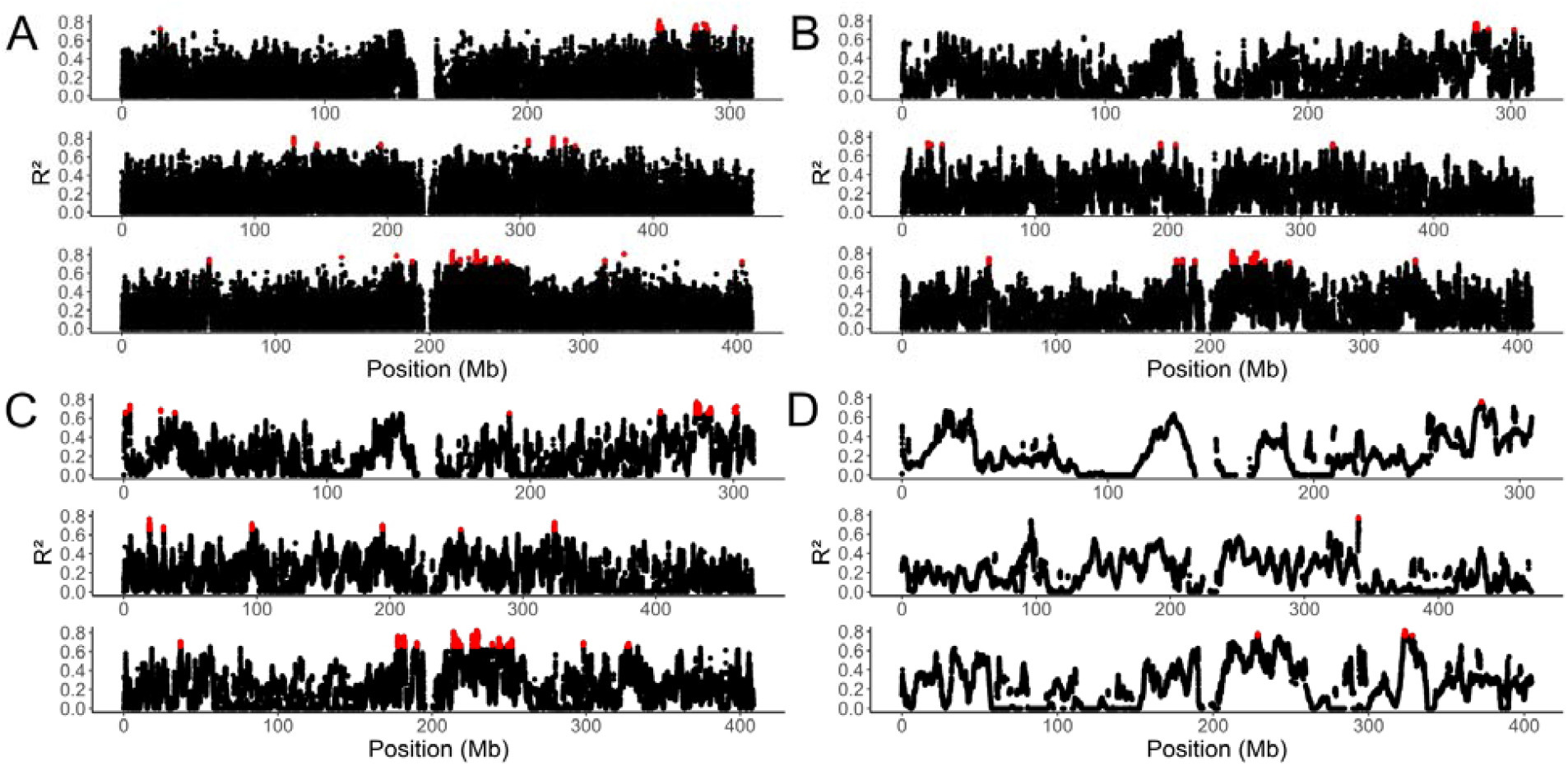
Predictive peaks from f4-ratio analysis is robust to changing window size. Manhattan plots showing the R^2^ from a GAM run at each individual window, red denotes windows that crossed the percentile that maximized predictive power for varying window sizes A) 100kb B) 500kb C) 1mb and D) 5mb.

